# Heparin flexibility within the extracellular matrix determines the bioactivity of bound vascular endothelial growth factor

**DOI:** 10.1101/2025.03.11.642311

**Authors:** Giuseppe Trapani, Daniele Di Iorio, Jurij Froese, Saskia Winter, Seraphine V. Wegner, Kay Grobe, Britta Trappmann

**Author notes:** <u>Correspondence:</u> Prof. Dr. Britta Trappmann.

## Abstract

Vascular endothelial growth factor (VEGF), a major regulator of blood vessel formation, is naturally bound to heparan sulfate proteoglycans in the extracellular matrix (ECM). Yet, how the physical presentation of VEGF by the matrix impacts its signaling potential remains fully unknown. To address this question, we have developed a tunable heparin-containing hydrogel model that recapitulates natural VEGF binding modes with full and independent control over physical properties. Using this model, we show that the degree of heparin flexibility in the hydrogel network determines the mobility of bound VEGF and, in turn, its ability to interact with VEGF receptor 2. We demonstrate that VEGF mobility is driven by a relay mechanism, in which VEGF molecules directly switch from one heparin chain to another. When strong electrostatic interactions between heparin and other hydrogel constituents immobilize the sugar backbone, this relay mechanism is impeded, in turn reducing VEGF’s ability to reach its target receptor at the cell membrane, thereby reducing its bioactivity. This work identifies heparin flexibility within the ECM as a previously unknown regulator of the microenvironment, which will not only contribute to a better mechanistic understanding of how the ECM regulates growth factor bioactivity, but it will also provide an important design criterion for the development of tissue-engineered biomaterials that require vascularization.

## 1. Introduction

The extracellular matrix (ECM) is a major regulator of the tissue microenvironment, as it presents resident cells with immobilized biomolecules that activate a variety of cell membrane receptors. Here, the physical matrix presentation of some of these biomolecules has emerged as a major regulator of their signaling potential; in particular, adhesive ligands are known to strongly activate integrins only when displayed on stiff matrices^1,2^. However, if and how the physical presentation of many other naturally matrix-bound biomolecules, in particular growth factors, regulates their activity, is still largely unknown.

Vascular endothelial growth factor-A_165_ (here referred to as VEGF) is a key regulator of angiogenesis during health and disease^3^. For example, VEGF signaling has emerged as a main driver of tumor progression, as it initiates and guides the growth of new blood vessels emanating from existing vasculature of the surrounding healthy tissue towards the tumor^4^. Due to its importance, VEGF signaling has been extensively studied to date; yet, its full mechanism of action remains elusive. While most approaches have studied the role of soluble VEGF, the growth factor is naturally bound to heparan sulfate proteoglycans (HSPGs) in the ECM^5,6^. Indeed, recent studies have suggested that this matrix binding itself alters the signaling potential, but details into the underlying molecular mechanisms are totally lacking^7^. Literature reports speculate that VEGF immobilization to matrices that are sufficiently stiff to activate β1 integrins prolongs the activity of its transmembrane receptor, VEGF receptor 2 (VEGFR-2), eliciting sustained downstream signaling^7^. However, anchoring to stiff matrices also affects the VEGF physical presentation, potentially impairing growth factor mobility and, in turn, its ability to physically interact with and elicit VEGFR-2 activation. Given this discrepancy, it is essential to understand how the physical presentation of VEGF regulates its signaling potential. Yet, this understanding is currently lacking due to the absence of suitable model matrices that recapitulate natural VEGF binding modalities, while allowing independent control over physical matrix properties, such as stiffness or the flexibility of bound biomolecules.

To fill this gap, synthetic hydrogels mimicking natural ECMs with full and independent control over the physical presentation of different types of bioactive molecules, in addition to the overall matrix stiffness, could serve as ideal model substrates^8,9^. While previous studies have already functionalized synthetic poly(ethylene glycol) (PEG)-based hydrogels with heparin^10^, an easily accessible and commercially available natural polymer that recapitulates an important natural VEGF binding site of HSPGs in tissues^11,12^, independent control over physical matrix properties, in particular heparin flexibility, is lacking in these systems.

Here, we develop a dextran-based hydrogel which combines the growth factor capturing ability of heparin with the finely tunable mechanical properties of a synthetic hydrogel. We report that heparin chain flexibility within the matrix network determines VEGF mobility, thereby regulating the bioactivity of the matrix-bound growth factor.

## 2. Results

### Design of matrix-tethered VEGF bioactivity assay

To study how VEGF matrix tethering regulates its signaling potential, we developed a sandwich assay in which endothelial cells were presented with synthetic hydrogel surfaces that offer precise control over VEGF nanoscale presentation (Fig. 1a, b). The synthetic hydrogels were based on a backbone of methacrylated dextran (DexMA)^13,14^ and methacrylated heparin (HepMA) (DexHepMA, 9:1 DexMA:HepMA) (Fig. 1c and Supplementary Fig. 1), the latter enabling electrostatic binding of VEGF similar to natural tissue matrices^5,12^. Since previous studies have shown that VEGFR-2 co-localizes with integrins on the cell membrane and that efficient VEGF signaling requires integrin crosstalk^15,16^, the cysteine-functionalized adhesive peptide RGD was coupled to the DexMA polymer backbone by Michael-type addition to methacrylate moieties. Gelation was induced by crosslinking of methacrylated polymer chains with matrix metalloproteinase (MMP)-sensitive bis-cysteine peptides (Fig. 1a), mimicking the degradable nature of natural ECMs. Changes in crosslinker content gave access to hydrogels of varying stiffness^13^, thereby presenting a tool to tune the physical presentation of the matrix-bound growth factor. After hydrogel formation, the surfaces were functionalized with VEGF molecules, which bind to the highly negatively charged polysaccharide chains of heparin^17,18^ with high affinity. To determine the level of bioactivity of matrix-bound VEGF, hydrogels were placed atop a monolayer of human umbilical vein endothelial cells (HUVECs) (Fig. 1b), which are known to respond to soluble VEGF by phosphorylation of the transmembrane receptor VEGFR-2 (Supplementary Fig. 2), resulting in activation of downstream signaling^19^. After treatment, VEGFR-2 activation was probed by determining its phosphorylation levels using Western blotting. Indeed, a short (2 min) contact with DexHepMA hydrogels functionalized with 330 ng/mL VEGF resulted in substantial levels of VEGFR-2 phosphorylation in HUVECs (Fig. 1d, e). This response was specific to the binding of VEGF to heparin, as DexMA hydrogels without HepMA, but incubated with VEGF, did not elicit any detectable activation of VEGFR-2 (Fig. 1d, e). Additionally, VEGFR-2 phosphorylation required covalent integration of heparin into the hydrogel network, since non-methacrylated heparin did not evoke any response (Supplementary Fig. 3). Together, these experiments demonstrate the suitability of the developed hydrogel-based assay to study the regulation of VEGF signaling by the physical presentation of the ECM.

**Fig. 1.**
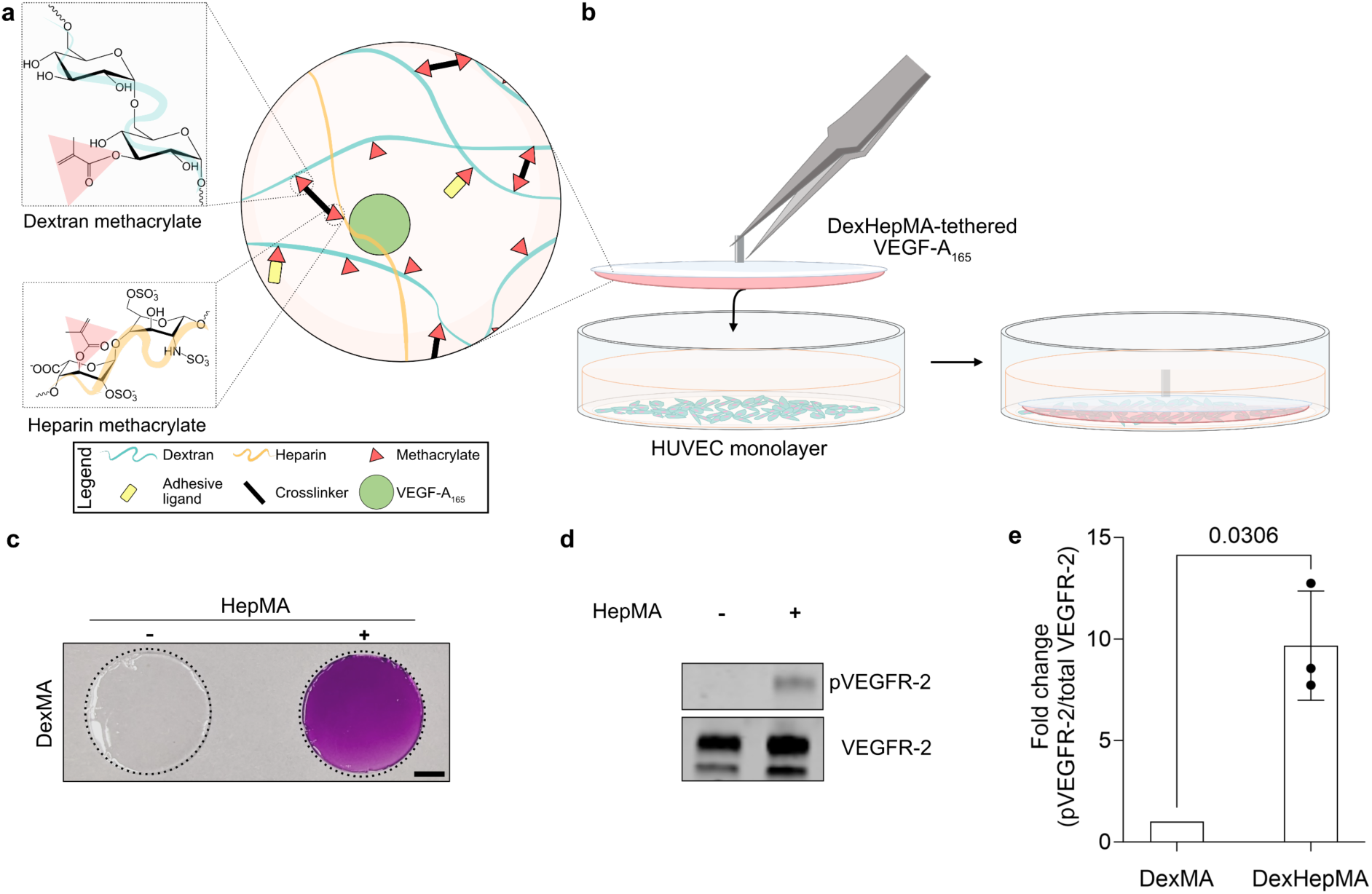
Design of an *in vitro* assay to assess matrix-tethered VEGF bioactivity. **a** Schematic representation of a DexHepMA hydrogel. A mixture of DexMA (functionalized with the cell-adhesive ligand CGRGDS) and HepMA polymer chains are crosslinked with MMP-cleavable peptides. VEGF binds to the hydrogel surface through electrostatic interactions with the negatively charged HepMA. **b** Sandwich assay set-up. VEGF-functionalized hydrogels are brought in contact with a HUVEC monolayer for the required assay time. Then, the hydrogel is removed and cells are lysed for further analysis. **c** Images of dimethylmethylene blue (DMMB)-stained hydrogels (outlined by dotted lines) with (right) and without (left) HepMA. Scale bar, 3 mm. **d** Activation of VEGFR-2 in HUVECs treated with VEGF-tethered hydrogels. HUVEC monolayers were exposed to VEGF-functionalized (330 ng/mL) hydrogels with (+) and without (-) HepMA containing 26 mM crosslinker for 2 min, followed by immunoprecipitation of VEGFR-2 and Western blot analysis for pan-phosphorylation of tyrosine residues (using 4G10 antibody) (top panel). As loading control, the blots were reprobed with an antibody against total human VEGFR-2 (bottom panel). **e** Quantification of fold change of pVEGFR-2 signal intensities, adjusted to the level of precipitated VEGFR-2 and normalized to HUVEC sample treated with VEGF-functionalized (330 ng/mL) hydrogels without HepMA. All data are reported as mean ± SD, p < 0.05 is considered to be statistically significant (two-tailed unpaired t-test with Welch’s correction) (n = 3 independent experiments).

### Matrix stiffness influences tethered VEGF bioactivity

Next, we investigated how changes in the physical presentation of matrix-tethered VEGF modulates its signaling potential. Specifically, we presented cells with VEGF tethered to hydrogels of varying stiffness (Fig. 2a, b), a major matrix cue that changes across different tissue environments and during the onset and progression of many diseases^20^. We hypothesized that changes in matrix stiffness could impact the bioactivity of tethered VEGF because of changes in the flexibility of the polymer network that VEGF is bound to. Interestingly, when VEGF was bound to lightly crosslinked/soft (26 mM crosslinker, ∼1 kPa) DexHepMA hydrogels, high levels of VEGFR-2 phosphorylation were observed, while VEGF bound to highly crosslinked/stiff (43.4 mM crosslinker, ∼5.9 kPa) hydrogels elicited >3.5-fold lower activation of VEGFR-2 (Fig. 2c, d), despite similar total VEGF surface concentrations across the two samples (Supplementary Fig. 4). To investigate whether these differences in VEGF matrix presentation impact downstream signaling as well, we analyzed the activity of the mitogen-activated protein kinase (MAPK) signaling pathway which is triggered by VEGFR-2 activation and which promotes, amongst many other cellular functions, cell proliferation^19^. In line with our previous observation, we detected a ∼2-fold decrease of extracellular-signal-regulated kinase (ERK) 1 and 2 phosphorylation levels when HUVECs were presented with VEGF bound to a stiff DexHepMA hydrogel, compared to a soft matrix (Fig. 2e, f). This observation suggests that matrix stiffness influences matrix-tethered growth factor bioactivity.

**Fig. 2.**
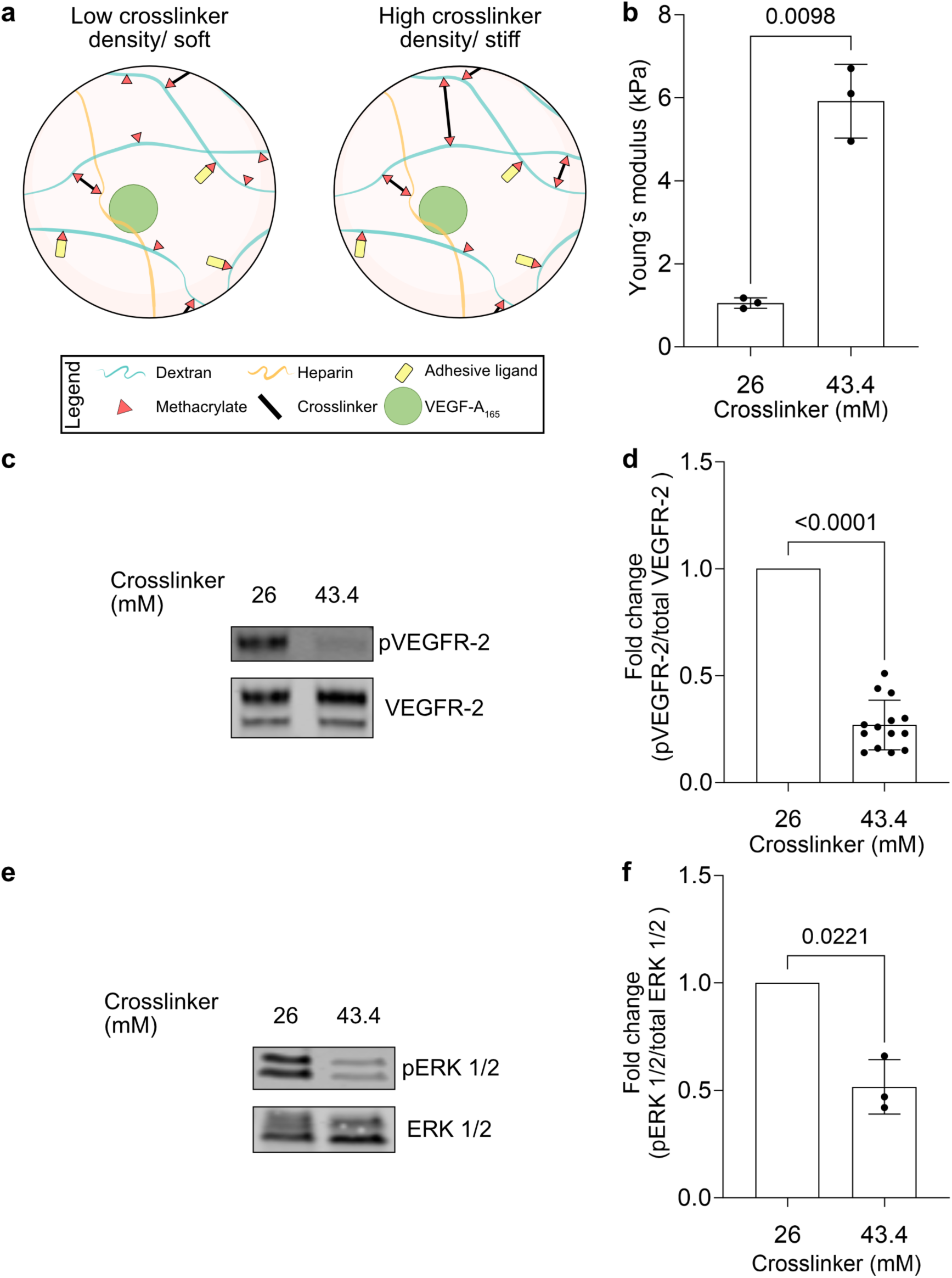
Signaling of tethered VEGF is influenced by matrix stiffness. **a** Design of DexHepMA hydrogels with tunable stiffness (determined by concentration of MMP-cleavable crosslinker). **b** Young’s moduli of DexHepMA hydrogels as a function of crosslinker concentration (n = 3 independent samples). **c** HUVEC monolayers were exposed to VEGF-functionalized (330 ng/mL) DexHepMA hydrogels with varying crosslinker concentrations for 2 min, followed by immunoprecipitation of VEGFR-2 and Western blot analysis for pan-phosphorylation of tyrosine residues (using 4G10 antibody) (top panel). As loading control, the blots were reprobed with an antibody against total human VEGFR-2 (bottom panel). **d** Quantification of fold change of pVEGFR-2 signal intensities, adjusted to the level of precipitated VEGFR-2 and normalized to HUVEC sample treated with VEGF-functionalized (330 ng/mL) DexHepMA hydrogel containing 26 mM crosslinker (n = 14 independent experiments). **e** HUVEC monolayers were exposed to VEGF-functionalized (330 ng/mL) DexHepMA hydrogels with varying concentrations of crosslinker for 15 min, followed by Western blot analysis of cell lysate using an antibody against human phospho-ERK 1/2. As loading control, the blots were reprobed with an antibody against total human ERK 1/2 (bottom panel). **f** Quantification of fold change of phospho-ERK 1/2 signal intensities, adjusted to the level of ERK 1/2 and normalized to HUVEC sample treated with VEGF-functionalized (330 ng/mL) DexHepMA hydrogel containing 26 mM crosslinker (n = 3 independent experiments). All data are reported as mean ± SD, p < 0.05 is considered to be statistically significant (two-tailed unpaired t-test with Welch’s correction).

### Matrix positive charge density impacts VEGF bioactivity

In our assay, VEGF is electrostatically bound to heparin molecules, and hence, its bioactivity is likely not only determined by the overall polymer network flexibility (i.e. hydrogel stiffness), but also by the flexibility of the heparin chains, which is a direct function of how tightly they are integrated in the matrix network. Here, one parameter to consider is the highly negatively charged nature of heparin molecules, which undergo electrostatic interactions with positively charged matrix constituents nearby, directly affecting the flexibility of heparin chains. Indeed, our hydrogels (similar to many other literature-known synthetic hydrogel systems) are crosslinked with positively charged peptides that carry a net charge of +1.91 (at pH 7, see Supplementary Table 1). Consequently, variations in hydrogel crosslinking to control stiffness are also accompanied by differences in the positive charge density of the matrix (Fig. 3a, Supplementary Table 2), and hence, heparin chain flexibility. Specifically, the highly crosslinked DexHepMA hydrogel (43.4 mM crosslinker) is not only stiffer than the lightly crosslinked hydrogel (26 mM crosslinker), but it is also characterized by a 1.7-fold increase in positive matrix charge density. We therefore entertained an alternative hypothesis for the effect of matrix stiffness on tethered VEGF bioactivity (Fig. 2): In highly crosslinked/stiff matrices with increased density of positive charges, stronger electrostatic interactions with negatively charged heparin chains reduce their flexibility, which could possibly impact the bioactivity of bound VEGF. To discern between the relative contributions of matrix stiffness and matrix positive charge density to the differences in tethered VEGF bioactivity, we increased heparin chain flexibility independently of changes in matrix stiffness by shielding the electrostatic interactions between heparin and the crosslinker peptide with the negatively charged polymer poly-L-glutamic acid (PGA) (Fig. 3b and Supplementary Fig. 5). Strikingly, presentation of VEGF bound to a highly crosslinked/stiff DexHepMA hydrogel, which would generally not induce VEGFR-2 phosphorylation, elicited >2-fold higher receptor activation when PGA was incorporated, thereby rescuing VEGF bioactivity on stiff matrices (Fig. 3c, d). This observation is in line with our hypothesis that increased heparin chain flexibility, through weaker electrostatic interactions between heparin and the hydrogel network, promotes VEGF bioactivity.

**Fig. 3.**
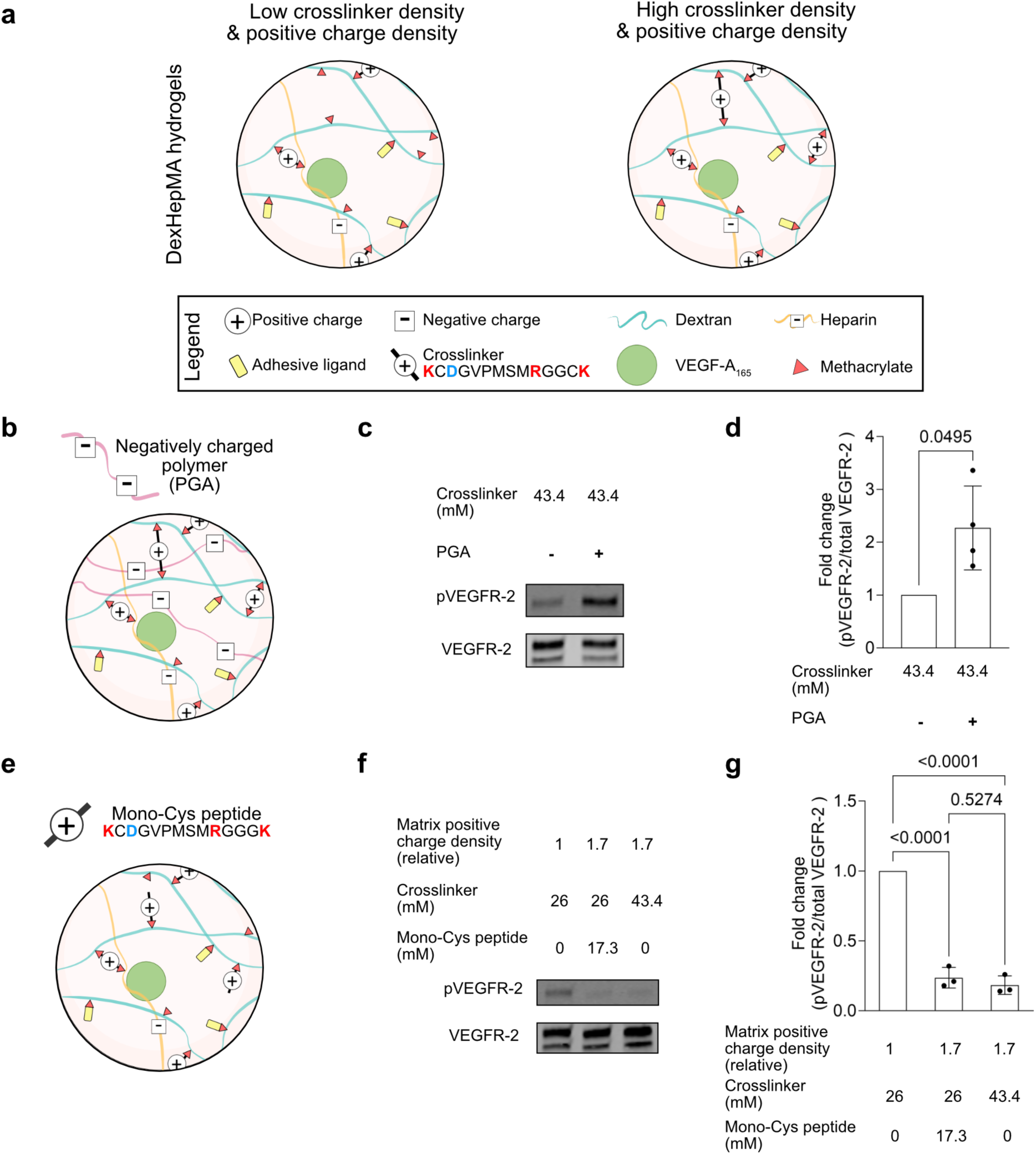
Nanoscale flexibility of heparin impacts signaling of matrix-bound VEGF. **a** Design of DexHepMA hydrogels with control over matrix positive charge density and crosslinking. **b** Schematic representation of a stiff DexHepMA hydrogel with embedded negatively charged PGA shielding peptide crosslinker positive charges. **c** VEGF-functionalized (330 ng/mL) DexHepMA hydrogels with (+) or without (-) PGA (85.3 mM) and crosslinked with 43.4 mM peptide crosslinker were presented to HUVEC monolayers for 2 min, followed by immunoprecipitation of VEGFR-2 and Western blot analysis for pan-phosphorylation of tyrosine residues (using 4G10 antibody) (top panel). As loading control, the blots were reprobed with an antibody against total human VEGFR-2 (bottom panel). **d** Quantification of fold change of pVEGFR-2 signal intensities, adjusted to the level of precipitated VEGFR-2 and normalized to HUVEC sample treated with VEGF-functionalized (330 ng/mL) DexHepMA hydrogel containing 43.4 mM crosslinker without PGA (n = 4 independent experiments). **e** Schematic representation of a soft DexHepMA hydrogel functionalized with mono-Cys peptide to match the positive charge density of a stiff hydrogel. **f** VEGF-functionalized (330 ng/mL) DexHepMA hydrogels of varying positive charges densities (value reported relative to the positive charge density of DexHepMA hydrogel containing 26 mM crosslinker without mono-Cys peptide) were presented to HUVEC monolayers for 2 min, and analysed as described in **c**. **g** Quantification of fold change of pVEGFR-2 signal intensities, adjusted to the level of precipitated VEGFR-2 and normalized to HUVEC sample treated with VEGF-functionalized (330 ng/mL) DexHepMA hydrogel containing 26 mM crosslinker without mono-Cys peptide (n = 3 independent experiments). All data are reported as mean ± SD, p < 0.05 is considered to be statistically significant. (One-way ANOVA with Tukeýs post-hoc test performed in **g**, two-tailed unpaired t-test with Welch’s correction in **d**).

While the introduction of PGA not only impacts the charge density of the matrix, but also the chemical composition of the hydrogel, we could not rule out any side-effects of chemical property changes on VEGF bioactivity. Therefore, rather than introducing an additional polymer in our hydrogel network, we next tuned the total density of positive charges through coupling a mono-cysteine variant of the charged crosslinker peptide (mono-Cys peptide) as a pendant chain to the hydrogel, thereby keeping the chemical composition as well as hydrogel stiffness constant (Fig. 3e, Supplementary Fig. 5). Specifically, we coupled the pendant mono-Cys peptide to the soft (∼1 kPa) DexHepMA hydrogel (Fig. 3e) to match the positive charge density of the stiff (∼5.9 kPa) hydrogel condition without concurrently affecting its stiffness (Supplementary Table 2, Supplementary Fig. 5). Remarkably, when VEGF was presented by soft, but strongly positively charged DexHepMA hydrogels functionalized with the mono-Cys peptide, phosphorylation levels of VEGFR-2 decreased ∼4-fold, compared to less positively charged hydrogels of the same stiffness without the mono-Cys peptide (Fig. 3f, g). This result shows that in the absence of any other confounding parameters, the positive charge density of matrices, rather than their stiffness, regulates the bioactivity of tethered VEGF. Together, our findings demonstrate the dependence of VEGF bioactivity on the flexibility of its binding partner heparin, which in turn is a result of electrostatic interactions within the matrix network.

### Heparin flexibility impacts VEGF relay between heparin chains

We next wanted to understand the molecular mechanism by which changes in the degree of heparin flexibility regulate the bioactivity of bound VEGF. Since we observed a drastic increase in cell surface receptor phosphorylation when VEGF was bound to more flexible heparin chains, we hypothesized that the degree of heparin chain flexibility changes the mobility of bound VEGF molecules, thereby enhancing their ability to reach the cell membrane receptor. To test this hypothesis, we reduced the complexity of our hydrogel system to gain independent control over the nanoscale flexibility of heparin chains. Specifically, we designed a 2D surface model solely based on the interactions between the charged heparin molecules and crosslinker peptides, which determine heparin nanoscale flexibility in our hydrogels. In order to mimic the hydrogel scenario, in which heparin molecules are not statically fixed at a rigid surface, but instead possess a high degree of freedom due to the flexibility of the overall polymer network, we based this model on supported lipid bilayers (SLBs) functionalized with heparin molecules, such that they can together move in 2D on an underlying glass support, and to which VEGF was bound electrostatically^21^. To test whether changes in heparin flexibility impact VEGF mobility, we incubated the heparin monolayer with the positively charged crosslinker peptide (20 mg/mL) prior to VEGF binding, in order to establish similar electrostatic interactions experienced by heparin chains within the hydrogel network. Using fluorescence recovery after photobleaching (FRAP), we directly visualized the mobility of heparin-bound fluorescein isothiocyanate (FITC)-labeled VEGF molecules in the absence or presence of crosslinker (Fig. 4a, b). Strikingly, in presence of the positively charged crosslinker peptide, we observed a 30% decrease in the FITC-labeled VEGF mobile fraction compared to the control without peptide (Fig. 4b). This result clearly shows that the level of heparin flexibility impacts the mobility of VEGF. Transferred to our hydrogel system, it confirms that heparin flexibility, which is a function of how tightly heparin chains are bound to the polymer network, determines the mobility of bound VEGF.

**Fig. 4.**
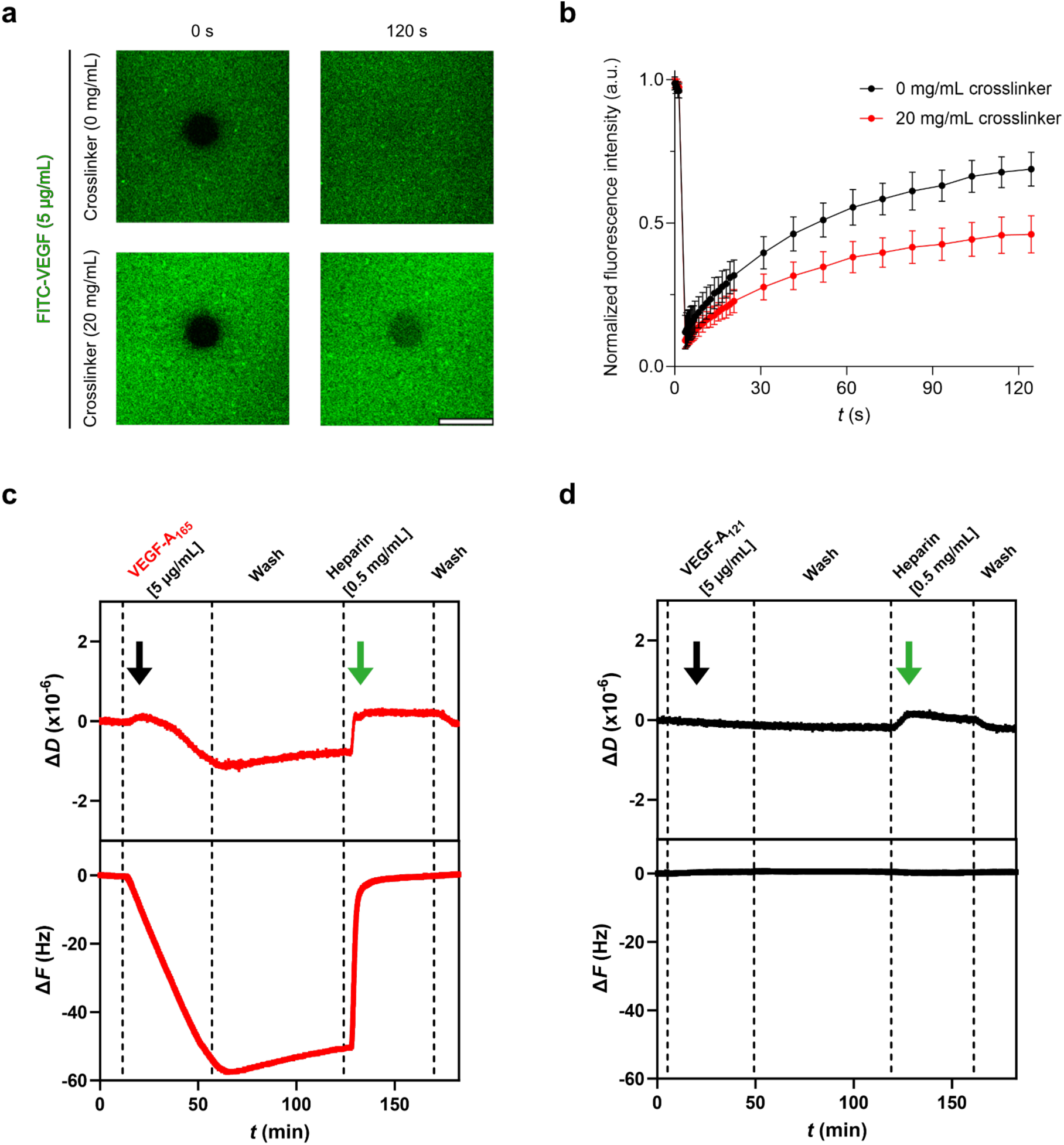
Heparin chain flexibility regulates VEGF mobility. **a** FRAP of FITC-labeled VEGF protein (5 µg/mL) interacting with heparin-functionalized SLBs with (top) or without (bottom) positively charged peptide crosslinker (20 mg/mL). Representative confocal images of FRAP assay, showing areas before (0 s) and after photo-bleaching (120 s). Scale bar, 20 µm. **b** Fluorescence intensity recovery profiles of FRAP experiments shown in (**a**) (n ≥ 3 analyzed bleached areas, pooled from 3 independent experiments). **c, d** Representative QCM-D binding assays of different VEGF isoforms (**c**, VEGF-A_165_; **d**, VEGF-A_121_) (5 µg/mL) interacting with heparin-functionalized SLBs. After protein interaction (black arrow) and buffer washing (PBS), soluble heparin (0.5 mg/mL) (green arrow) was flushed into the QCM-D chamber to compete with SLB-bound heparin chains.

To induce VEGFR-2 phosphorylation, VEGF needs to physically associate with its receptor at the cell membrane. Our sandwich and FRAP assays together demonstrate that, in samples with less flexible heparin chains, the limited mobility of matrix-tethered VEGF impedes its interaction with VEGFR-2. This observation suggests that VEGF mobility within the matrix is driven by a mechanism that requires heparin chain flexibility. Indeed, we recently proposed a model of protein transport between highly sulfated polysaccharide chains for the morphogen Hedgehog (Hh), whose mobility is determined by its ability to simultaneously bind two chains (chain crosslinking) and switch between them^21^, meaning that the physical proximity between the protein and the sugar chain likely determines the degree of protein mobility. In light of this model, we hypothesized that, similar to Hh, VEGF mobility in the matrix relies on heparin chain cross-over, in turn regulating its ability to reach VEGFR-2 at the cell membrane. In fact, VEGF molecules are homodimers with two heparin-binding domains^12^, and thus fulfill the essential requirement of binding two heparin chains simultaneously in order to move directly from one chain to another. To test this hypothesis, we performed quartz crystal microbalance with dissipation monitoring (QCM-D) experiments on heparin-functionalized SLBs (Fig. 4c, d), which allow to characterize real-time mass variations as a function of changes in resonance frequency (Δ*F*), whereby VEGF binding to heparin results in a decrease in Δ*F*, while increased Δ*F* indicates protein unbinding. Additionally, variations in energy dissipation (Δ*D*) can be correlated with changes in the viscoelasticity of the heparin monolayer, where a decrease in Δ*D* indicates layer rigidification^21^. As shown in Fig. 4c, VEGF-A_165_ isoform (5 µg/mL) incubation with the SLB induced a decrease in frequency, - Δ*F*, indicating VEGF-A_165_ binding to heparin (Fig. 4c, black arrow). This variation in frequency was accompanied by a proportional decrease in dissipation, -Δ*D*, indicative of a stiffening of the heparin layer consistent with VEGF-mediated heparin chain crosslinking. Notably, VEGF remained bound to the heparin surface even during extensive washing, pointing out a low unbinding rate (off-rate constant) of the protein. In contrast, after injection of soluble heparin (0.5 mg/mL) onto the VEGF-loaded heparin layer, VEGF-A_165_ immediately detached from the QCM-D chip (Fig. 4c, green arrow). Here, soluble heparin acted as an acceptor of VEGF, allowing for the protein to relay from one chain to another, accelerating its elution from the sensor surface^21^. To demonstrate that the differences observed were specific to captured VEGF, we repeated the QCM-D experiments with VEGF-A_121_, an isoform that lacks the ECM binding site and is known to be unable to interact with heparin^6^, where the dissipation and frequency remained unchanged throughout the treatments (Fig. 4d). Together, the results of the QCM-D experiments clearly demonstrate that, similarly to Hh, heparin-associated VEGF-A_165_ moves by switching between chains, corroborating our hypothesis that the flexibility of heparin chains within the hydrogel network regulate changes in VEGF bioactivity by impacting this relay mechanism.

### Off-rate for VEGF-heparin binding impacts VEGF mobility

While we have demonstrated that VEGF movement within the matrix is influenced by the flexibility of heparin chains, the mobility of tethered growth factors is also likely affected by the ability of one binding domain to dissociate from the matrix, which determines how easily the growth factor transitions to a neighboring heparin chain. We therefore hypothesized that, in addition to heparin chain mobility, VEGF bioactivity is influenced by the off-rate of its binding to heparin. Specifically, lower off-rates for VEGF-heparin binding would hinder the relay of the growth factor, preventing VEGF from reaching and physically interacting with its target receptor on the cell membrane. To investigate this hypothesis, we tethered a different VEGF-A isoform, VEGF-A_189_, which exhibits a lower off-rate for ECM HSPGs^6,22^.

First, we compared the mobility of VEGF-A_165_ and -A_189_ isoforms on an SLB-supported heparin layer by QCM-D. Interestingly, compared to VEGF-A_165_ (Fig. 5a), a solution of VEGF-A_189_ isoform induced a stronger decrease in dissipation (Fig. 5b, black arrow). This observation is best shown by the Δ*D*/-Δ*F* plot, where, for each unit of Δ*F*, VEGF-A_189_ causes a more pronounced variation in Δ*D*, compared to VEGF-A_165_ (Fig. 5c). This marked variation in dissipation indicates a more rigid heparin layer in presence of VEGF-A_189_ resulting from enhanced crosslinking of the sugar chains. This observation was further supported by flushing soluble heparin (0.5 mg/mL) into the QCM-D chamber, as VEGF-A_189_ switched from the sensor surface to soluble heparin acceptors more slowly than VEGF-A_165_ (Fig. 5a, b, green arrow). Together, the QCM-D experiments clearly show that heparin-bound VEGF-A_189_ is characterized by lower mobility from one heparin chain to the next and hence, slower movement within the matrix. Subsequently, we tested in the sandwich assay whether this diminished mobility of matrix-bound VEGF-A_189_ isoform determined a loss of protein bioactivity. Indeed, when HUVECs were exposed to VEGF-A_189_ bound to both soft or stiff DexHepMA hydrogels, we detected low levels of VEGFR-2 phosphorylation (Fig. 5d, e), even though cells were clearly responsive to soluble VEGF-A_189_ (Supplementary Fig. 6). This result demonstrates that increased VEGF-heparin interactions result in a loss of VEGF bioactivity. Together, our findings establish the dependence of VEGF bioactivity on its nanoscale mobility, which is impacted by the flexibility of its binding partner heparin, as well as its affinity to the matrix.

**Fig. 5.**
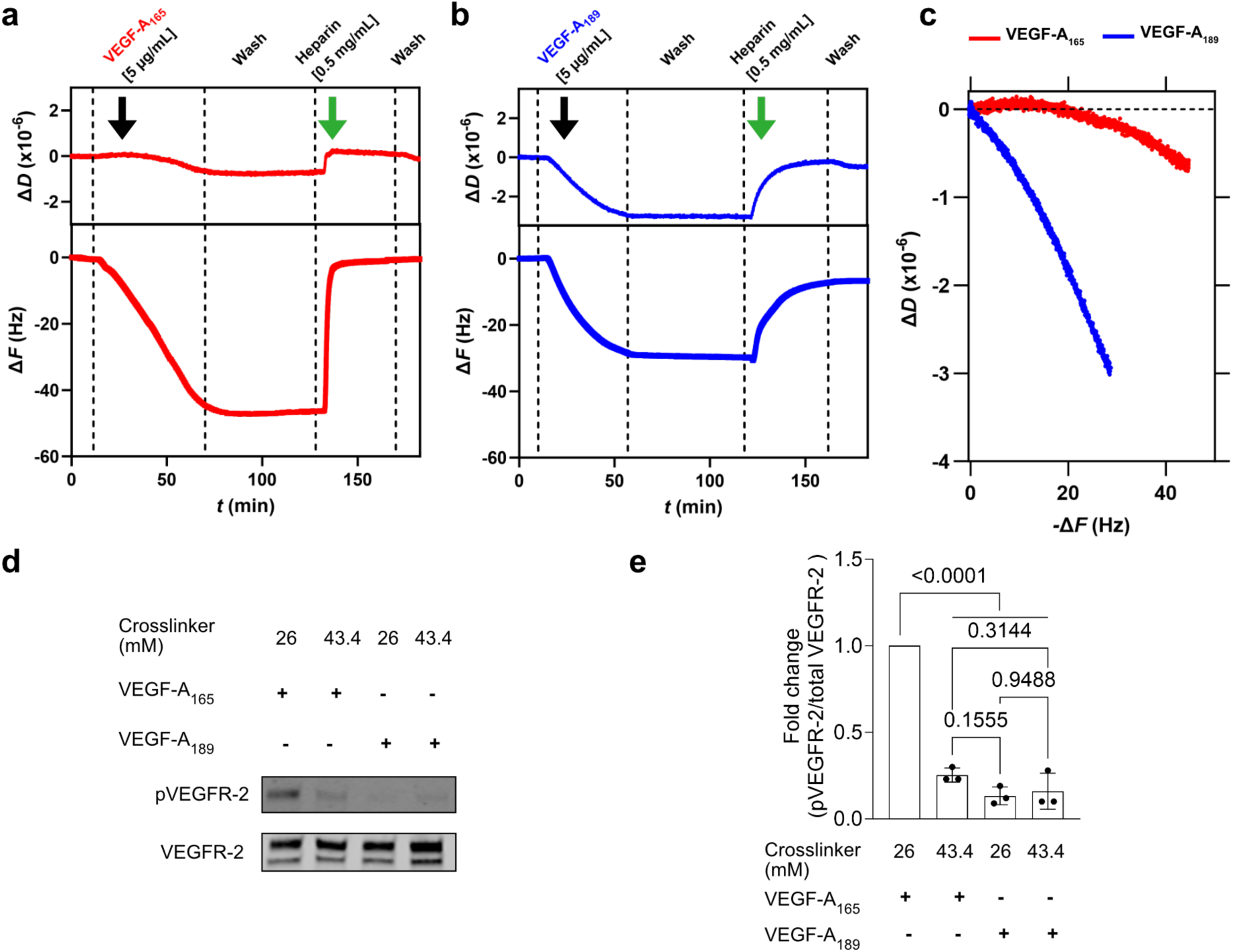
Off-rate for VEGF-heparin binding determines growth factor mobility. **a**, **b** Representative QCM-D binding assays of different VEGF isoforms (a, VEGF-A_165_; b, VEGF-A_189_) (5 µg/mL) interacting with heparin-functionalized SLBs. After protein interaction (black arrow) and buffer washing (PBS), soluble heparin (0.5 mg/mL) (green arrow) was flushed into the QCM-D chamber to compete with SLB-bound heparin chains. **c** Δ*D* as a function of -Δ*F* for the VEGF isoforms (red, VEGF-A_165_; blue, VEGF-A_189_) derived from the protein interaction portion of the QCM-D plots (black arrow) reported in (**a**, **b**). **d** VEGF-A_165_ and VEGF-A_189_-functionalized (330 ng/mL) DexHepMA hydrogels with varying concentrations of positively charged peptide crosslinker were presented to HUVEC monolayers for 2 min, followed by immunoprecipitation of VEGFR-2 and Western blot analysis for pan-phosphorylation of tyrosine residues (using 4G10 antibody) (top panel). As loading control, the blots were reprobed with an antibody against total human VEGFR-2 (bottom panel). **e** Quantification of fold change of pVEGFR-2 signal intensities, adjusted to the level of precipitated VEGFR-2 and normalized to HUVEC sample treated with VEGF-A_165_-functionalized (330 ng/mL) DexHepMA hydrogel containing 26 mM of peptide crosslinker (n = 3 independent experiments). All data are reported as mean ± SD, p < 0.05 is considered to be statistically significant (one-way ANOVA with Tukeýs post-hoc test).

### Impact of VEGF mobility on cell function

Having established the dependence of matrix-bound VEGF bioactivity on its mobility in assays that probed immediately occurring signaling events, we next wanted to know if differences in VEGF matrix presentation also have long lasting functional consequences for endothelial cells. In particular, we thought to study whether proliferation and cytoskeletal remodeling, two major processes involved in VEGF-driven angiogenesis, are continuously impacted by longer-term culture on matrices with different VEGF mobilities. In order to do so, we directly cultured HUVECs on the surface of VEGF-functionalized DexHepMA hydrogels with varying degrees of heparin flexibility. Instead of our standard stiffnesses that were used for the sandwich assay, we chose hydrogels that were extremely soft (17.4 mM crosslinker, ∼0.3 kPa) (Supplementary Fig. 5), since we wanted to minimize the impact of integrin activation on proliferation and cytoskeletal remodeling^23^ as a confounding parameter. VEGF mobility was changed by introduction of mono-Cys peptide (25.9 mM) to increase the density of positive matrix charges (matching a stiff ∼5.9 kPa hydrogel) (Supplementary Table 2), which in our previous experiments resulted in reduced bioactivity due to low mobility of VEGF. We found that although cell adhesion and spreading occurred both in presence and absence of mono-Cys peptide (due to the presence of the adhesive ligand RGD), striking differences in cell aspect ratio occurred (Fig. 6a, b).

**Fig. 6.**
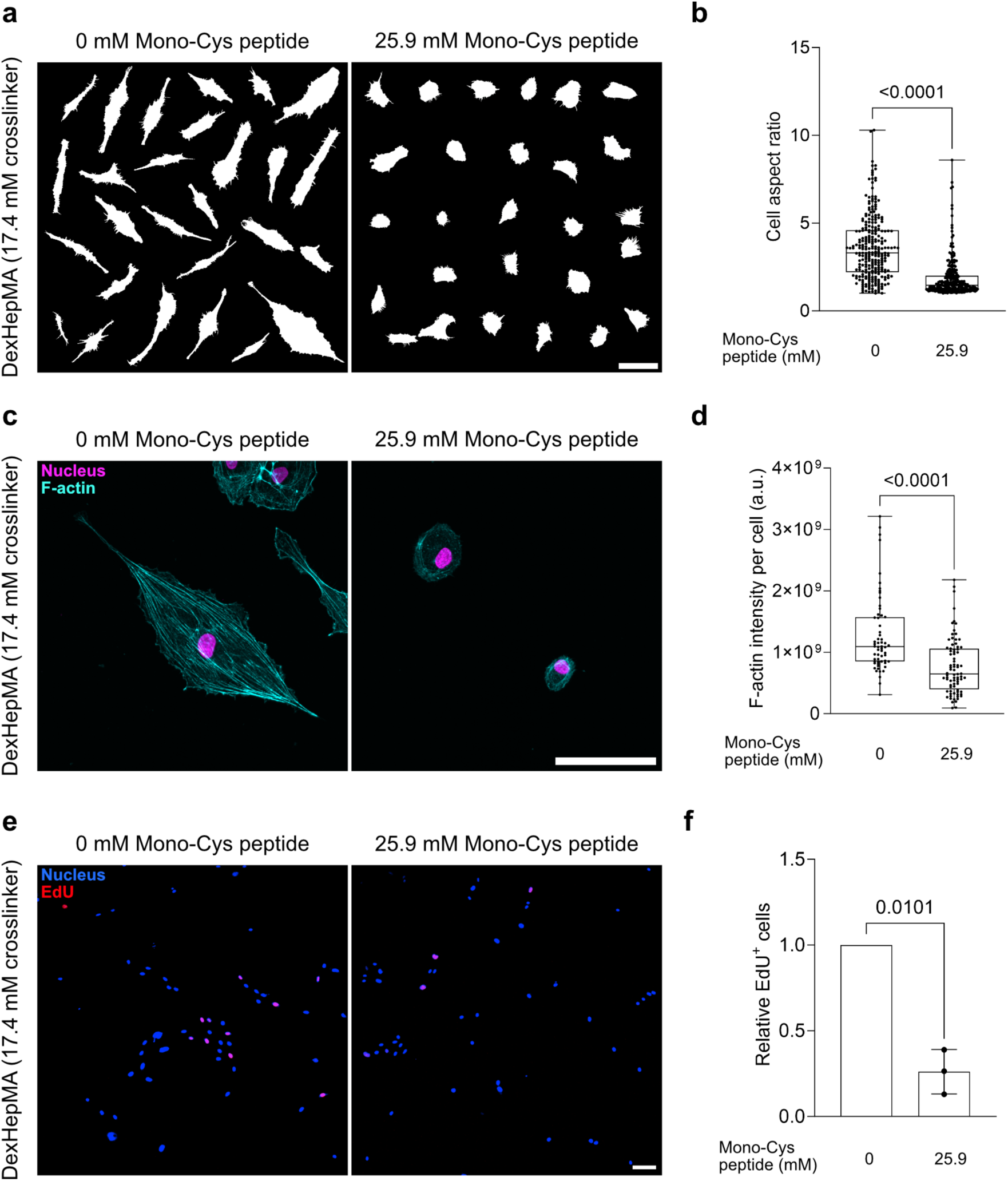
Heparin chain flexibility regulates VEGF-induced remodeling of the cellular actin cytoskeleton and cell proliferation rates. **a** Representative cell outlines of HUVECs cultured atop VEGF-functionalized (330 ng/mL) DexHepMA hydrogels, with (right) or without (left) 25.9 mM mono-Cys peptide, crosslinked with 17.4 mM crosslinker peptide. **b** Quantification of cell aspect ratio for conditions shown in (**a**) (n > 200 cells, pooled from 3 independent experiments). **c** Representative high magnification confocal images of HUVECs on hydrogels described in (**a**). Composite fluorescence images show nuclei (magenta) and F-actin (cyan). **d** Quantification of F-actin fluorescence intensity per cell after overnight culture (> 16 h) on hydrogels described in (**a**) (n > 50 cells, pooled from 3 independent experiments). **e** Representative confocal images of HUVECs after overnight culture (> 16 h) on hydrogels described in (**a**) and treated for 4 h with EdU. Composite fluorescence images show nuclei (blue) and EdU (red). **f** Quantification of EdU^+^ cells for conditions shown in (**e**). The number of EdU^+^ cells were normalized to the number of EdU^+^ HUVECs cultured on VEGF-functionalized (330 ng/mL) DexHepMA hydrogels crosslinked with 17.4 mM crosslinker peptide and containing 0 mM mono-Cys peptide (n = 3 independent experiments). Data in (**b**) and (**d**) are reported as box-and-whiskers plots (box, 25–75 percentile; bar-in-box, median; whiskers, the minimum and maximum values); data in (**f**) are represented as mean ± SD, p < 0.05 is considered to be statistically significant (two-tailed unpaired t-test with Welch’s correction). Scale bar, 100 µm.

Specifically, on DexHepMA hydrogels with higher VEGF nanoscale mobility (condition without mono-Cys peptide), HUVECs acquired a more elongated cell shape, whereas cells remained mostly round on hydrogels with reduced VEGF mobility (condition with mono-Cys peptide). The same elongated phenotype was acquired by HUVECs seeded on control DexHepMA hydrogels (without mono-Cys peptide, not functionalized with VEGF) treated with the soluble form of the growth factor (Supplementary Fig. 7), demonstrating that the observed morphological difference is a direct consequence of VEGF signaling. The elongated phenotype, which was only observed when VEGF was presented by matrices with more heparin chain flexibility, is reminiscent of a pro-migratory phenotype, which has been shown to be regulated by VEGFR-2^24^, indicating that indeed, VEGF signaling was efficiently activated in these conditions only. In addition to cell shape, we also observed a profound remodeling of the endothelial actin cytoskeleton towards more prominent stress fibers when HUVECs were cultured atop DexHepMA hydrogels with more mobile VEGF (condition without mono-Cys peptide) (Fig. 6c). This observation was accompanied by an increase in the total amount of F-actin present in the cell (∼1.7-fold increase compared to condition with mono-Cys peptide) and increased assembly of focal adhesions (Fig. 6c, d and Supplementary Fig. 8), in line with literature reports on VEGF-induced endothelial cell actin cytoskeleton remodeling^25^. In addition to the actin cytoskeleton, we tested the effect of differences in VEGF matrix-binding on HUVEC proliferation rates (Fig. 6e, f). In line with the observed decrease in ERK 1/2 phosphorylation in HUVECs treated with VEGF bound to less flexible, stiff hydrogels in the short-term sandwich assay (Fig. 2e, f), the number of cells that had proliferated over a culture time of 4 h had dropped >3.5-fold in the presence of mono-Cys peptide (determined by EdU assay) (Fig. 6e, f). Hence, it is clear that changes in VEGF mobility not only regulate short-term cell processes, but greatly control essential cellular functions during longer-term tissue culture.

Taken together, our studies highlight the role of the matrix in regulating the signaling potential of bound VEGF. Based on our findings, we developed a model (Fig. 7) that describes how heparin flexibility, governed by electrostatic interactions with the surrounding hydrogel network, determines the mobility of bound VEGF, and its potential to reach the target receptor at the cell membrane: in lightly crosslinked/soft matrices, weaker electrostatic interactions between the negatively charged heparin chains and the positively charged hydrogel crosslinker render heparin chains more flexible and therefore allow VEGF to easily switch from one chain to another, reaching and activating VEGFR-2 at the cell surface (curved arrows represent VEGF relay, Fig. 7, left panel). In contrast, in highly crosslinked/stiff matrices with higher positive charge densities, heparin chains are more tightly immobilized within the hydrogel, diminishing tethered-VEGF mobility, in turn, preventing its engagement with VEGFR-2 (Fig. 7, right panel). This model, for the first time, uncovers physical ECM properties as a major regulator of VEGF bioactivity.

**Fig. 7.**
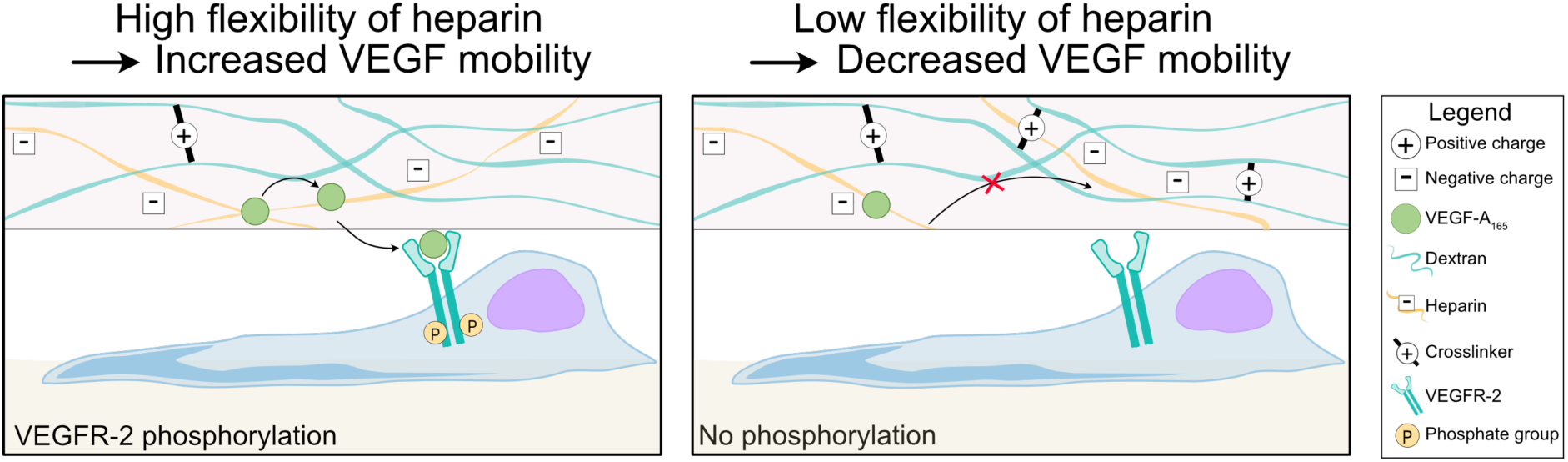
Model of heparin flexibility regulating matrix-tethered VEGF mobility and, in turn, its bioactivity. (Left) In lightly crosslinked/soft hydrogels, fewer electrostatic interactions between heparin and the surrounding matrix (in our case positive crosslinker charges) lead to increased flexibility of heparin chains, compared to stiff matrices. This heparin flexibility increase promotes matrix-bound VEGF relay (curved arrows) required to reach VEGFR-2 at the cell surface and to induce receptor phosphorylation. (Right) In a stiff/highly crosslinked hydrogel, the more positively charged matrix electrostatically immobilizes heparin chains within the hydrogel network and decrease their flexibility. In turn, matrix-bound VEGF molecules are unable to relay from one heparin chain to another, decreasing their overall mobility, and hampering their interactions with VEGFR-2, which results in decreased receptor phosphorylation.

## 3. Discussion

VEGF has long been established as a major microenvironmental regulator of blood vessel formation and function, and hence, the molecular mechanisms underlying its signaling have been investigated in great detail^26^. While the vast majority of findings are based on the addition of soluble VEGF, VEGF in natural tissues is bound to the ECM^5^; the possibility that the mode of physical VEGF presentation by the matrix alone impacts growth factor bioavailability and -activity has largely been overlooked to date. Using a synthetic hydrogel, we found that the mobility of VEGF, which is impacted by the flexibility of its binding partner heparin, is a major regulator of growth factor bioactivity.

The impact of matrix binding on VEGF signaling has been the focus of several published reports^7,27,28^. For example, Chen et al. found that VEGF binding to collagen type I hydrogels promotes VEGFR-2 association with β1 integrin, altering receptor tyrosine phosphorylation patterns and prolonging VEGFR-2 activation^7^. However, since collagen is an unnatural VEGF binding partner, it is unclear if and how these mechanistic insights can be transferred to more physiological ECMs. To recapitulate native VEGF binding modalities, recent studies developed heparin-incorporated hydrogel models^10^, mimicking the high affinity binding of VEGF to HSPGs^5,11^; how the physical presentation of heparin by the matrix regulates the bioactivity of bound VEGF, has not been investigated or considered. The study that comes closest to addressing the impact of VEGF matrix presentation has bound VEGF to heparin-functionalized gold surfaces to show that covalent locking of the growth factor orientation has surprisingly little impact on VEGFR-2 activation^27,28^. While this finding suggests that the VEGF nanoscale presentation is not a major regulator of its signaling potential, the study lacked to investigate the role of heparin flexibility, and how this impacts the mobility of bound VEGF. In our work, by incorporating heparin within a mechanically tunable synthetic hydrogel that recapitulates native VEGF binding modalities, we found that the flexibility of heparin within the hydrogel network influences the mobility of bound VEGF molecules by regulating their ability to switch between heparin chains and, therefore, to reach their target receptor at the cell membrane.

In this study, we report the importance of heparin flexibility within the ECM network in the regulation of growth factor bioactivity. To the best of our knowledge, this is the first example that a structural ECM component, which captures and presents bioactive molecules, plays an active role in regulating cell function through its physical matrix presentation. Instead, previous studies have mainly focused on the importance of the physical matrix presentation of bioactive signaling molecules whose activity is regulated by integrin-mediated mechanotransduction, such as mechanoresponsive growth factors or adhesive ligands ^29–31^. For example, similar to our observations on heparin flexibility, pioneering work by Hubbell and co-workers has shown that efficient integrin activation and cell spreading requires the adhesive peptide RGD to be coupled to PEG diacrylate hydrogels through a flexible spacer arm in order to ensure accessibility of the otherwise sterically unavailable ligand^32^. Our study shows that the importance of physical ECM presentation is not exclusive to molecules that directly signal to cells, but extends to components that capture and present bioactive molecules, such as HSPGs. One major limitation which has previously impeded the investigations in this direction is the lack of model materials that allow to decouple the effect of heparin flexibility from co-changing parameters, e.g. matrix stiffness. In our study, we fill this gap by our development of a synthetic DexHepMA hydrogel model that offers independent control over the nanoscale flexibility of heparin molecules through changing their network integration, in addition to matrix stiffness.

Mobility of matrix-stored VEGF has classically been thought to occur through diffusion resulting from association to/ dissociation from ECM components^22,33^. If VEGF mobility was indeed primarily driven by diffusion, then differences would only result from changes in the strength of interaction with its ECM binding partner, which is quantified by its association/dissociation constants^18,34^. However, in our key experiment, we keep the affinity of the heparin/VEGF pair constant, and only change heparin flexibility; yet, we see a major change in VEGF mobility. This finding calls for an alternative model of VEGF mobility that is not based on diffusion. Here, we demonstrate that matrix-bound VEGF mobility is driven by an intersegmental growth factor transfer between heparin chains. This transfer is supported by the two binding sites that are present in the VEGF-A_165_ structure, a direct consequence of its dimeric nature with one heparin-binding domain in each monomer^6,12^. We speculate that the relay mechanism is initiated as one VEGF domain dissociates from one heparin molecule to switch to a nearby chain, while the second domain still remains bound. This process repeats for both domains, thereby driving VEGF movement. Compared to free 3D diffusion, this movement is a lot more efficient, as VEGF follows a pre-determined, pseudo-2D path. This efficiency enhancement is conceptually analogous to certain well-studied intracellular transport phenomena, such as the binding and movement dynamics between DNA-binding proteins and DNA^35,36^. Importantly, our model is in line with our previous work, which had for the first time shown a similar relay mechanism for HSPG-binding partners, in particular the morphogen Hh and the homotrimeric spike protein of SARS-CoV-2^21,37^. Here, we extend this mechanism to other biomolecules which bind reversibly to HSPGs and demonstrate the profound impact the mode of ECM binding has on biomolecule activity.

Our finding that VEGF needs to be presented by heparin molecules that are flexibly incorporated in ECMs in order to retain its bioactivity has major implications for the field of tissue engineering, which has over the last decade made tremendous efforts to develop implantable biomaterials that promote vascularization^38–40^. Historically, soluble VEGF has been incorporated into implant materials^41^; however, a major limitation is its rapid clearing from the material target site, not leaving sufficient time for the endothelial cells from the surrounding healthy vasculature to form new blood vessels towards the implant. To overcome this hurdle, more recent advances have coupled VEGF to biomaterials^42,43^, thereby leveraging the restricted diffusivity and spatial control over the tethered VEGF to initiate angiogenesis-related signaling events. With our work, we now establish VEGF presentation through heparin matrix integration as an important design criterion that needs to be taken into account for the design of vascularized biomaterials.

To summarize, our studies establish VEGF flexibility as a previously unknown cue that regulates growth factor bioactivity. While in the current study, we have used synthetic hydrogels due to their high level of control over matrix nanoscale properties, the findings will be highly relevant for natural tissue ECMs which are composed of many different types and concentrations of charged biomolecules. Importantly, the ECM composition not only varies between different tissue types at homeostasis, but also during the progression of many diseases in which matrix remodeling plays a major role^44^. Based on our observations, changes in growth factor flexibility resulting from differences in matrix charges could be a major regulator of these processes. In addition to a better mechanistic understanding of VEGF signaling, our findings will also aid in the design of tissue engineered constructs that rely on vascularization. Here, future studies will have to determine the impact of VEGF flexibility on blood vessel formation in 3D. Our developed hydrogel model will be directly applicable to this question since its MMP cleavable crosslinks allow for 3D cell and tissue culture applications. In addition to VEGF signaling, our well-controlled hydrogel models will be broadly applicable to study other growth factor families that are naturally matrix-bound and which are likely impacted by their nanoscale matrix presentation as well.

## 5. Experimental Section

### Reagents

All reagents were purchased from Sigma Aldrich, unless otherwise stated.

### Cell-adhesive and MMP-cleavable peptide sequences

The cell-adhesive peptide CGRGDS and the MMP-cleavable peptide sequences KCDGVPMSMRGGCK and KCDGVPMSMRGGGK (mono-Cys peptide) were custom synthesized (provided as HCl salt) by GenScript Biotech at >95% purity.

### Antibodies

Monoclonal mouse anti-phosphotyrosine 4G10 antibody was purchased from Merck Chemicals (#05-321). Polyclonal rabbit anti-human phospho-p44/42 MAPK (ERK1/ERK2) (Thr 202, Tyr 204) antibody (#36-8800) and Alexa Fluor™ 647 conjugated donkey anti-mouse secondary antibody (#A31571) were purchased from Thermo Fischer Scientific. Monoclonal rabbit anti-human VEGFR-2 (55B11) (#2479) and monoclonal mouse anti-human p44/42 MAPK (ERK1/ERK2) (L34F12) (#4696) antibodies were purchased from Cell Signaling Technology. IRDye® 800CW donkey anti-mouse IgG (#926-32212), IRDye® 800CW donkey anti-rabbit IgG (#926-32213), IRDye® 680RD donkey anti-rabbit IgG (#926-68073) and IRDye® 680RD donkey anti-mouse IgG (#926-68072) secondary antibodies were purchased from LI-COR. Mouse monoclonal anti-vinculin antibody was purchased from Sigma Aldrich (#V9131).

### Synthesis of DexMA

Dextran methacrylation was performed following a previously published procedure^14,45^. Briefly, dextran (20 g, MW 86 000 Da, MP Biomedicals) and 4-dimethylaminopyridine (2 g, Fluka Analytical) were dissolved in anhydrous dimethyl sulfoxide (100 mL). Glycidyl methacrylate (24.6 mL) was added and the mixture was vigorously stirred for 24 h at 45 °C in the dark. Next, the solution was transferred on ice and precipitated into ice cold 2-propanol (1 L, VWR chemicals). The precipitated product was retrieved by centrifugation and solubilized in Milli-Q^®^ water. The solution was purified by dialysis (SnakeSkin™ dialysis Tubing, Life Technologies, 10 kDa MW cutoff) against Milli-Q^®^ water for three days with two water exchanges daily. Finally, purified DexMA was obtained by lyophilization and characterized by ^1^H-NMR spectroscopy using deuterium oxide (D_2_O) as solvent. A 0.7 methacrylate/dextran repeat unit ratio was determined.

### Synthesis of HepMA

HepMA was prepared by reacting heparin sodium salt with methacrylic anhydride. In brief, methacrylic anhydride (450 µL) was added to a 1% w/v heparin (100 mg, from porcine intestinal mucosa, Sigma-Aldrich #H3149) solution in PBS (10 mL) and reacted for 4 h in an ice bath while maintaining the pH at 7.5–8.5 with 1 M NaOH. Then, fresh methacrylic anhydride (450 µL) was added and the reaction allowed to proceed for an additional 4 h while maintaining the pH at 7.5–8.5. The reaction mixture was kept under vigorous stirring overnight at room temperature. Subsequently, the aqueous phase was dialyzed (SnakeSkin™ dialysis Tubing, Life Technologies, 3.5 kDa MW cutoff) against Milli-Q^®^ water for six days. Then, the solution was sterile filtered (Filtropur S Syringe filter, Sarstedt, 0.2 µm pore size), lyophilized and the final product was characterized by ^1^H-NMR spectroscopy using D_2_O as solvent. The degree of heparin methacrylation was calculated as the ratio between the integral of the vinyl protons (5.77 and 6.21 ppm) and the anomeric protons of the heparin disaccharide unit (5.24 and 5.43 ppm). A methacrylate/heparin disaccharide repeat unit ratio of 0.06 was determined.

### Preparation of DexHepMA hydrogels for growth factor tethering

For the preparation of DexHepMA hydrogels, mixtures of DexMA (final concentration of 4% w/v) and the adhesive peptide CGRGDS (final concentration of 6 mM) were prepared in Dulbeccós modified Eagle medium (DMEM) (pH 7.0, without sodium bicarbonate) (Gibco™), the pH was adjusted to 8 using 1 M NaOH and the reaction allowed to proceed for 30 min. Next, a mixture of HepMA (final concentration 0.4% w/v) and KCDGVPMSMRGGCK crosslinker (final concentration 17.4, 26 or 43.4 mM), dissolved in DMEM pH 7.0, were added to the reaction solution, the pH was readjusted to 8 and the hydrogel precursor solution was allowed to polymerize for 30 min at room temperature. For the preparation of different DexHepMA hydrogel formulations, the following compounds were introduced in the reaction mixture after the addition of the crosslinker: KCDGVPMSMRGGGK mono-Cys peptide (final concentration 25.9 or 17.3 mM) and PGA (poly-L-glutamic acid sodium salt, MW 15-50 kDa, Sigma-Aldrich #P4761) (final concentration of the repeating unit 85.3 mM), dissolved in DMEM pH 7.0, for the preparation of DexHepMA hydrogel with mono-Cys and DexHepMA hydrogel with PGA, respectively. For the preparation of DexMA hydrogels deprived of HepMA, the protocol was modified to increase the concentration of DexMA to 4.4% w/v; to introduce heparin, a solution of heparin sodium salt (final concentration 0.4% w/v), in DMEM pH 7.0, was added after the crosslinker (final concentration 26 mM) addition.

### Casting of flat hydrogels

To ensure good contact between the hydrogel and the HUVEC monolayer during cell treatments, flat hydrogel surfaces were generated by polymerization of the precursor solution between two 25 mm round cover glasses. The bottom glass (hydrogel recipient) surface was not treated and therefore sticky to the DexHepMA hydrogel, it was also equipped with magnets to allow precise positioning of the hydrogels on the HUVEC monolayer using tweezers. The top surface (used for flat casting) was functionalized with a sacrificial gelatin gel which after melting left a separation between the hydrogel and the glass surface to ensure easy coverslip removal. Specifically, the hydrogel precursor solution (80 µL) was cast on a round cover glass (VWR), with an attached magnet (3 mm diameter x 6 mm height, magnets4you GmbH) (glued e.g. with cured polydimethylsiloxane (PDMS) (Dow Chemical Company)) on the outer side, and flattened by placing on top of it a gelatin-functionalized round cover glass of the same size. Briefly, gelatin-functionalized round cover glasses were prepared by incubating plasma treated cover glasses with a 2% v/v (3-aminopropyl)triethoxysilane (APTES) solution in ethanol overnight at room temperature; cover glasses were incubated for 1 h at room temperature with glutaraldehyde, then washed, sterilized and functionalized with a 1% w/v gelatin solution overnight at 37 °C. Before usage, gelatin-functionalized cover glasses were extensively washed in water and sonicated. After polymerization, the gelatin-functionalized cover glass was removed and the hydrogel was washed in PBS overnight at 4 °C.

### Growth factor functionalization of DexHepMA hydrogels

Hydrogels were washed twice with 1 M NaCl solution and twice with PBS at 4 °C, to remove non-covalently bound HepMA or heparin polymer chains. Next, VEGF was tethered to the hydrogels by incubating with 300 µL solution of 330 ng/mL recombinant human VEGF-A_165_ and VEGF-A_189_ (R&D Systems) in PBS for 2 h at room temperature. Subsequently, hydrogels were washed and stored in PBS overnight at 4 °C. DexHepMA hydrogels with PGA (and respective PGA-free controls in the same experiment) were washed four times with PBS only (instead of NaCl) to avoid the detachment of PGA polymer chains from the hydrogel network due to high salt concentrations. Hydrogels used for cell treatments and non-VEGF functionalized control gels were simply incubated with PBS following the same number of washing steps reported above.

### DMMB hydrogel staining

DMMB staining solution was prepared following a previously published report^46^. Briefly, DMMB (16 mg) was dissolved in an aqueous solution (1 L, pH 3) containing glycine (3.04 g), NaCl (2.37 g) and HCl (0.1 M, 95 mL). DexMA and DexHepMA hydrogels, washed as described above, were incubated overnight in DMMB solution at room temperature in the dark on a platform rocker. Then, hydrogels were extensively washed with PBS at room temperature in the dark to remove any unbound dye molecule.

### Cell culture

HUVECs pre-screened for VEGF response were obtained from PromoCell^®^ (#C-12205). Cells were cultured in fully supplemented endothelial cell growth medium 2 (EGM-2) (PromoCell^®^) containing 250 ng/mL amphotericin B and 10 μg/mL gentamicin (Gibco™). For sandwich assay experiments, freshly trypsinized HUVECs were resuspended in EGM-2 and seeded on 25 mm round Nunc™ Thermanox™ plastic coverslips (Thermo Fischer Scientific), placed in a 6-well plate, at a density of 18,300 cells/cm^2^ and cultivated for 3 days. For cell morphology and proliferation studies, freshly trypsinized HUVECs were resuspended in EGM-2 (without or supplemented with 30 ng/mL VEGF) and seeded on DexHepMA hydrogels (with or without mono-Cys peptide) at a density of 4,200 cells/cm^2^ and cultivated overnight (> 16 h). Passage 5 cells were used for all experiments.

### Matrix-tethered growth factor activity assay (sandwich assay)

Prior to the assay, confluent HUVECs were starved overnight in endothelial cell basal medium 2 (EBM-2) (PromoCell^®^) containing 250 ng/mL amphotericin B, 10 μg/mL gentamicin (Gibco™) and 0.5% fetal calf serum (FCS). Cells were presented with the VEGF-functionalized hydrogels for varying time points (2 min for VEGFR-2 activation, 15 min for ERK activation) by inverting them on the monolayer. For soluble VEGF treatment controls, cells were incubated with 30 ng/mL VEGF-A_165_ or VEGF-A_189_ solution in EBM-2. Negative controls were treated with EBM-2 only. To ensure good contact between the cell monolayer and the hydrogel surface, tissue culture dishes containing the HUVEC monolayer were placed on a metal plate to attract the magnet attached to the cover glass supporting the hydrogel.

### Western blot analysis

After treatment, cell cultures were placed on ice. Whenever HUVEC monolayers were treated with tethered VEGF surfaces, the hydrogels were peeled off carefully. HUVECs were rinsed twice with cold PBS (with Ca^2+^ and Mg^2+^) (Corning) and lysed in lysis buffer (20 mM Tris-HCl, pH 7.4, 150 mM NaCl, 2 mM CaCl_2_, 1% Triton X-100, 0.1 % NaN_3_, 20 µM Na_3_VO_4_, 1 cOmplete™ Mini EDTA-free Proteinase Inhibitor Cocktail (Roche)) for 5 min on ice. Lysates were further incubated on a revolver rotator for 20 min at 4°C, and centrifuged at 16,100 g for 30 min at 4°C. Solutions of comparable protein levels, determined using Pierce™ BCA protein assay kit (Thermo Scientific), were prepared by dilution with lysis buffer. For VEGFR-2 immunoprecipitation, 80 µL aliquots were incubated for 2 h at 4 °C on a tube revolver rotator with Protein A-Sepharose™ CL-4B (Cytiva™) loaded with anti-human VEGFR-2 antibody. Immunocomplexes were retrieved by centrifugation at 2300 g for 3 min at 4 °C and washed with lysis buffer five times. Total cell lysates and immunoprecipitated VEGFR-2 were diluted with SDS sample buffer (187.5 mM Tris–HCl pH 6.8, 200 mM dithiothreitol, 30% glycerol, 6% SDS, and 0.02% bromophenol blue) and heated to 95 °C for 5 min. Protein separation was performed by SDS-PAGE followed by wet blotting to transfer proteins to nitrocellulose membranes (Cytiva™). Membranes were washed with TBS-T (supplemented with 250 µM Na_3_VO_4_), and blocked for 1 h at room temperature in blocking buffer (TBS-T supplemented with 2% BSA). VEGFR-2 and ERK 1/2 phosphorylation levels were probed by anti-phosphotyrosine 4G10 (1 µg/mL in blocking buffer) and anti-human phospho-p44/42 MAPK (1:1000 dilution in blocking buffer) antibodies, respectively. VEGFR-2 and ERK 1/2 protein-loading was determined by anti-human VEGFR-2 (1:1000 in blocking buffer) and anti-human p44/42 MAPK (1:2000 in blocking buffer) antibodies, respectively. Primary antibodies were detected by fluorescently-labeled secondary antibodies (1:10000 in blocking buffer) and visualized by an Odyssey^®^ CLx imaging system (LI-COR^®^). Images were analysed by Image Studio Software (LI-COR^®^). Phosphorylated proteins were normalized to the respective non-phosphorylated proteins as loading controls.

### Hydrogel mechanical characterization by nanoindentation

Hydrogel Young’s moduli were measured by nanoindentation (Piuma, Optics 11 life, the Netherlands) using a cantilever with a 0.045 N/m spring constant and an attached bead of 48.5 µm diameter. Measurements were performed on thick hydrogels in PBS supplemented with 10% FCS at room temperature. Indentation curves were fitted with a Hertz contact model. At least 20 indentations from 3 independent samples were averaged for each hydrogel.

### Matrix-tethered VEGF quantification

Since VEGF was coupled to thick hydrogels in our studies, a significant proportion of VEGF molecules diffused into the 3D matrix and got trapped in its porous network, rendering quantification of surface-bound VEGF only with standard fluorescence staining procedures difficult. Moreover, the generally low Z-resolution of confocal microscopy, which was used for imaging, did not allow for a distinction between surface fluorescence signal and contributions from the bulk of the hydrogel. To overcome these limitations, we made use of a previously described microfluidic device^13,47^ in which a perfusable channel is embedded in a 3D hydrogel matrix, allowing for the visualization of the channel surface-bound growth factor in the x-y plane, in which confocal microscope resolution is highest. This setting closely mimicks the binding of VEGF to the hydrogel surface in the sandwich assay.

To quantify VEGF bound to DexHepMA hydrogels containing either 26 or 43.4 mM peptide crosslinker, FITC-labeled recombinant human VEGF-A_165_ was used. For protein labeling and purification, FluoReporter® FITC Protein Labeling Kit (Invitrogen) was used, following the manufactureŕs instructions. Briefly, recombinant human VEGF-A_165_ (Peprotech) was reacted with reactive dye (100:1 dye:protein monomer) in 0.1 M phosphate buffer (pH 7.4, 0.15 M NaCl) for 2 h at room temperature in the dark. Protein labeling reaction was followed by purification through spin columns provided by the labeling kit. A 0.3 degree of labeling (dye per protein dimer) was determined, assuming 85% recovery of labeled-protein. Purified FITC-labeled recombinant human VEGF-A_165_ was stored at –80 °C in 0.1 % BSA solution. For quantification of the channel surface-bound VEGF, a DexHepMA hydrogel containing either 26 or 43.4 mM peptide crosslinker was cast in the central chamber of the microfluidic device and, after polymerization, washed following the same washing steps reported above. FITC-labeled recombinant human VEGF-A_165_ was added to one channel of the microfluidic device at a concentration of 10 µg/mL in PBS and incubated for 2 h at room temperature on a platform rocker, PBS was added to the second, parallel channel. Then, devices were extensively washed and stored in PBS overnight at 4 °C on a platform rocker. Finally, FITC-labeled VEGF fluorescence signal was acquired.

### Synthesis of biotinylated heparin

Biotinylated heparin was synthesized by adapting a previously reported procedure^48^. In brief, heparin (heparin sodium salt BioChemica, PanReac AppliChem #A3004) was dissolved (final concentration 4 mM) in 100 mM acetate buffer (pH 4.5, prepared with glacial acetic acid (Carl Roth, Karlsruhe, Germany) and sodium acetate) containing aniline (100 mM). Biotin-PEG_3_-oxyamine (3.4 mM, Conju-Probe, San Diego, USA) was added to the heparin solution and allowed to react for 48 h at 37 °C. The solution was dialyzed against Milli-Q^®^ water for 48 h (3.5 kDa MW cutoff, Spectra/Por), and subsequently lyophilized for 48 h. For further use, the conjugates were diluted to the desired concentrations in wash buffer A (10 mM Tris, 100 mM NaCl at pH 7.4). The obtained biotinylated heparin was characterized by biotin-streptavidin binding assays using QCM-D.

### Preparation of SUVs

SUVs were prepared by adapting previously reported procedures^49,50^. A mixture of 1 mg/mL 1,2-dioleoyl-*sn*-glycero3-phosphocholine (DOPC, Avanti Polar Lipids) and 5 mol% of 1,2-dioleoyl-*sn*-glycero-3-phosphoethanolamine-*N*-(cap biotinyl) (DOPE-biotin, Avanti Polar Lipids) was prepared in chloroform in a glass vial. Subsequently, the solvent was evaporated by a low-flow nitrogen stream while constantly turning the vial to obtain a homogenous lipidic film. The residual solvent was removed for 1 h under vacuum. The dried film was rehydrated in Milli-Q^®^ to a final concentration of 1 mg/mL and vortexed to dissolve the lipidic film. The lipid dispersion was sonicated for about 15 min until the solution turned clear. The obtained SUVs were stored in the refrigerator and used within 2 weeks. For the FRAP experiments, lipid mixtures of DOPC and DOPE-biotin with 1,1′-dioctadecyl-3,3,3′,3′-Tetramethylindodicarbocyanine (DiD, Thermo-Fischer Scientific) (molar ratio 94.9: 5.0: 0.1) were used to form the SUVs.

### QCM-D measurements

QCM-D measurements were performed with a QSense Analyser (Biolin Scientific, Gothenburg, Sweden) and SiO_2_-coated sensors (QSX303, Biolin Scientific). The measurements were performed at 22 °C by using four parallel flow chambers and one peristaltic pump (Ismatec, Grevenbroich, Germany) with a flow rate of 75 µL per min. The normalized frequency shifts Δ*F*, and the dissipation shifts Δ*D*, were measured at six overtones (*i* = 3, 5, 7, 9, 11, 13). The fifth overtone (*i* = 5) was presented throughout; all other overtones gave qualitatively similar results. QCM-D sensors were first cleaned by immersion in a 2 wt% SDS solution and subsequent sonication for 10 min. Afterwards the QCM-D sensors were immersed in Milli-Q^®^, followed by sonication for 10 min. The sensors were then dried under a nitrogen stream and activated by 10 min treatment with a UV/ozone cleaner (Ossila, Sheffield, UK). For the formation of SLBs, after obtaining a stable baseline, freshly made SUVs were diluted to a concentration of 0.1 mg/mL in wash buffer A containing 10 mM of CaCl_2_ directly before use and flushed into the chambers. The quality of the SLBs was monitored in situ to ascertain that high-quality SLBs were formed, corresponding to equilibrium values of Δ*F* = −24 ± 1 Hz and Δ*D* < 0.5 × 10^−6^ ^51^. Afterwards, a solution of streptavidin (Sav; 150 nM, in wash buffer A) was flushed over the SLBs, followed by the addition of biotinylated heparin (10 μg per mL, in wash buffer A). Each sample solution was flushed over the QCM-D sensor until the signals were equilibrated, and subsequently rinsed with wash buffer A. Before the addition of VEGF protein solutions, the fully functionalized QCM-D sensors were equilibrated with PBS and the flow rate was reduced to 20 µL per min. The tested VEGF isoforms (VEGF-A_121_, VEGF-A_165_, and VEGF-A_189_) were diluted in PBS to a final concentration of 5µg/mL prior to flushing over the fully functionalized QCM-D sensors.

### FRAP measurements

For the FRAP experiments, SLBs were formed in 18-well microslides with a glass bottom (Ibidi, Gräfelfing, Germany), following a previously reported procedure^52^. First, wells were pre-treated with 150 μL of a solution of sodium hydroxide (2 M) for 1 h to form a hydrophilic surface. Next, the glass slides were rinsed three times with Milli-Q^®^ and three times with wash buffer A containing 10 mM CaCl_2_. Subsequently, a solution of freshly prepared SUVs (final concentration 0.1 mg/mL) was added to the wells containing buffer with 10 mM CaCl_2_ for 30 min at room temperature, leading to the formation of an SLB. Excess lipids were removed from the wells by rinsing with 100 μL buffer (without CaCl_2_) at least five times. Next, a solution of streptavidin (Thermo Fisher, final concentration 250 nM in wash buffer A) was added to the SLBs, followed by the addition of a solution of biotinylated heparin (final concentration 100 μg/mL in wash buffer A) and solutions of peptide crosslinker (final concentrations 20 mg/mL in PBS), which were incubated for 30 min each. Between each addition, wells were rinsed with buffer at least five times in order to rinse off the excess material. In particular, prior to peptide crosslinker addition, wells were extensively washed with PBS. Finally, a solution of FITC-labeled VEGF (final concentration 5 µg/mL in PBS) was added to the functionalized SLBs.

All FRAP measurements were performed with a Leica SP8 confocal laser scanning microscope through a 63 × water objective. A circular spot of ∼10 µm in diameter was bleached using a laser at 552 and 638 nm (100% intensity), and the recovery of the fluorescence intensity in the bleached regions was monitored. The FITC-labeled VEGF was excited with a 488 nm laser, and the emission was detected at 500 nm – 540 nm; the DiD dye was excited with a 638 nm laser, and the emission was detected at 655nm - 705nm. For both the SLB and the FITC-labeled VEGF, the FRAP protocol consisted of 5 imaging loops (0.37 s intervals) before bleaching, 5 loops during bleaching (0.37 s intervals), and 30 loops during recovery (10 at 0.37 s intervals, 10 at 1.37 intervals, and 10 at 10.37 s intervals). All images were analyzed by using Leica Application Suite X (LAS X) software version 3.7.1.21655.

### Fluorescent staining, microscopy, and image analysis

For cell proliferation and morphology studies, HUVEC cultures on glass coverslip-supported DexHepMA hydrogels were treated for 4 h with 10 µM EdU solution in EGM-2 without VEGF. Afterwards, cells were washed with PBS and fixed for 15 min with 4% paraformaldehyde (Thermo Fischer Scientific) at room temperature. Labeling of EdU^+^ cells was performed following the manufactureŕs protocol (Click-iT™ Plus EdU Alexa Fluor™ 555 Imaging Kit, Thermo Fischer Scientific). To visualize cell nuclei, fixed samples were incubated with Hoechst 33342 (Thermo Fisher Scientific, 1:250) in 0.1% Tween 20 solution in PBS at room temperature for 20 min. Samples were inverted and imaged by high-speed spinning disk confocal microscopy (Dragonfly by Andor with built-in software Fusion, v2.0.0.13) at 10× magnification. Quantification was performed by dividing the number of EdU^+^ cells by the total number of cells in each sample. Given the incompatibility of EdU ClickiT with phalloidin staining, visualization of the F-actin cytoskeleton was performed in a separate step after EdU visualization. Cells were incubated with Alexa Fluor™ 488 phalloidin (Thermo Fisher Scientific, 1:250) and Hoechst 33342 (1:250) in 0.1% Tween solution in PBS at room temperature overnight. Samples were inverted and imaged by confocal microscopy at 10× and 40× magnifications. Quantification of cell aspect ratio was performed for single cells using ImageJ software on maximum intensity projections of 10× images using the MinError autothresholding method. Quantification of F-actin intensity per cell was performed by ImageJ on the sum intensity projection of 40× images. The reported F-actin intensity per cell is the RawIntDen of F-actin (visualized by fluorescent phalloidin) measured within each individual cell outlined using the MinError autothresholding method. For focal adhesion visualization, samples were permeabilized in 0.5% Triton X-100 (Thermo Fisher Scientific) for 1 h, blocked with 3% BSA in PBS for 1 h and incubated with mouse monoclonal anti-vinculin antibody (1:100 in blocking buffer) for 1 h at room temperature. Then, samples were extensively washed with 0.1% Tween 20 in PBS, and incubated with a solution of Alexa Fluor™ 647 conjugated donkey anti-mouse antibody (1:500) for 1 h at room temperature. Prior to imaging, samples were extensively washed with 0.1% Tween 20 in PBS. Vinculin was imaged by confocal microscopy at 40× magnification. FITC-labeled VEGF intensity at the channel surface in the microfluidic device was defined as the highest intensity value of the plot profile acquired from the edge of channel towards the edge of the other, parallel channel. The highest fluorescence intensity was averaged over 25 measurements per image. All microscopic fluorescence images are presented as maximum intensity projections.

### Statistical Analysis

All statistical analyses were carried out using GraphPad Prism 9 (v 10.0.0). Statistical significance was determined by two-tailed unpaired Student’s t-test (with Welch’s correction where specified) or one-way ANOVA. p values < 0.05 were considered statistically significant. All data are presented as a mean ± standard deviation unless stated otherwise. Each study was independently repeated three times or as specified.

## Author contributions

G.T., K.G., S.V.W. and B.T. conceived the study, designed experiments, and interpreted results; G.T. conducted experiments and analyzed data; S.W. conducted cell treatments in sandwich assays and performed Western blotting; D.D.I. conducted FRAP experiments and analyses; J.F. conducted QCM-D experiments; G.T. and B.T. wrote the manuscript.

## Acknowledgements

The authors thank Dr. M. Pitulescu for constructive discussions and suggestions; Dr. S. Volkery, M. Stasch and Dr. N. Kirschnick (BioOptic Service unit, MPI Münster) for imaging support; U. Ipe for excellent technical support and the Department of Prof. Dr. D. Vestweber (MPI Münster) for giving access to equipment; Dr. K. Bergander and co-workers (NMR facility, Organic Chemistry Department of University of Münster) for giving access to the NMR spectrometer; Dr. M. Jain for constructive feedback on the manuscript. This work was financially supported by the Max Planck Society (MPG; B.T.) and the German Research Foundation (DFG, SFB1348: A07; B.T., A08; K.G., A14; S.V.W.). G.T. is supported by the International Max Planck Research School–Molecular Biomedicine, Münster, Germany.

## Data availability statement

The data that support the findings of this study are available from the corresponding author upon reasonable request.

## References

1 Yeung, T. et al. Effects of substrate stiffness on cell morphology, cytoskeletal structure, and adhesion. Cell Motil Cytoskeleton 60, 24–34 (2005). 10.1002/cm.20041

2 Discher, D. E., Janmey, P. & Wang, Y. L. Tissue cells feel and respond to the stiffness of their substrate. Science 310, 1139–1143 (2005). 10.1126/science.1116995

3 Apte, R. S., Chen, D. S. & Ferrara, N. VEGF in Signaling and Disease: Beyond Discovery and Development. Cell 176, 1248–1264 (2019). 10.1016/j.cell.2019.01.021

4 Weis, S. M. & Cheresh, D. A. Tumor angiogenesis: molecular pathways and therapeutic targets. Nat Med 17, 1359–1370 (2011). 10.1038/nm.2537

5 Hynes, R. O. The extracellular matrix: not just pretty fibrils. Science 326, 1216–1219 (2009). 10.1126/science.1176009

6 Houck, K. A., Leung, D. W., Rowland, A. M., Winer, J. & Ferrara, N. Dual regulation of vascular endothelial growth factor bioavailability by genetic and proteolytic mechanisms. J Biol Chem 267, 26031–26037 (1992).

7 Chen, T. T. et al. Anchorage of VEGF to the extracellular matrix conveys differential signaling responses to endothelial cells. J Cell Biol 188, 595–609 (2010). 10.1083/jcb.200906044

8 Lutolf, M. P. & Hubbell, J. A. Synthetic biomaterials as instructive extracellular microenvironments for morphogenesis in tissue engineering. Nat Biotechnol 23, 47–55 (2005). 10.1038/nbt1055

9 Trappmann, B. & Chen, C. S. How cells sense extracellular matrix stiffness: a material’s perspective. Curr Opin Biotechnol 24, 948–953 (2013). 10.1016/j.copbio.2013.03.020

10 Zieris, A. et al. FGF-2 and VEGF functionalization of starPEG-heparin hydrogels to modulate biomolecular and physical cues of angiogenesis. Biomaterials 31, 7985–7994 (2010). 10.1016/j.biomaterials.2010.07.021

11 Powell, A. K., Yates, E. A., Fernig, D. G. & Turnbull, J. E. Interactions of heparin/heparan sulfate with proteins: appraisal of structural factors and experimental approaches. Glycobiology 14, 17R–30R (2004). 10.1093/glycob/cwh051

12 Robinson, C. J., Mulloy, B., Gallagher, J. T. & Stringer, S. E. VEGF165-binding sites within heparan sulfate encompass two highly sulfated domains and can be liberated by K5 lyase. J Biol Chem 281, 1731–1740 (2006). 10.1074/jbc.M510760200

13 Trappmann, B. et al. Matrix degradability controls multicellularity of 3D cell migration. Nat Commun 8, 371 (2017). 10.1038/s41467-017-00418-6

14 van Dijk-Wolthuis, W. N. E. et al. Synthesis, Characterization, and Polymerization of Glycidyl Methacrylate Derivatized Dextran. Macromolecules 28, 6317–6322 (1995). 10.1021/ma00122a044

15 Soldi, R. et al. Role of alphavbeta3 integrin in the activation of vascular endothelial growth factor receptor-2. EMBO J 18, 882–892 (1999). 10.1093/emboj/18.4.882

16 Borges, E., Jan, Y. & Ruoslahti, E. Platelet-derived growth factor receptor beta and vascular endothelial growth factor receptor 2 bind to the beta 3 integrin through its extracellular domain. J Biol Chem 275, 39867–39873 (2000). 10.1074/jbc.M007040200

17 Fairbrother, W. J., Champe, M. A., Christinger, H. W., Keyt, B. A. & Starovasnik, M. A. Solution structure of the heparin-binding domain of vascular endothelial growth factor. Structure 6, 637–648 (1998). 10.1016/s0969-2126(98)00065-3

18 Zhao, W., McCallum, S. A., Xiao, Z., Zhang, F. & Linhardt, R. J. Binding affinities of vascular endothelial growth factor (VEGF) for heparin-derived oligosaccharides. Biosci Rep 32, 71–81 (2012). 10.1042/BSR20110077

19 Olsson, A. K., Dimberg, A., Kreuger, J. & Claesson-Welsh, L. VEGF receptor signalling - in control of vascular function. Nat Rev Mol Cell Biol 7, 359–371 (2006). 10.1038/nrm1911

20 Butcher, D. T., Alliston, T. & Weaver, V. M. A tense situation: forcing tumour progression. Nat Rev Cancer 9, 108–122 (2009). 10.1038/nrc2544

21 Gude, F. et al. Hedgehog is relayed through dynamic heparan sulfate interactions to shape its gradient. Nat Commun 14, 758 (2023). 10.1038/s41467-023-36450-y

22 Park, J. E., Keller, G. A. & Ferrara, N. Vascular Endothelial Growth-Factor (Vegf) Isoforms - Differential Deposition into the Subepithelial Extracellular-Matrix and Bioactivity of Extracellular Matrix-Bound Vegf. Mol Biol Cell 4, 1317–1326 (1993). DOI 10.1091/mbc.4.12.1317

23 Trappmann, B. et al. Extracellular-matrix tethering regulates stem-cell fate. Nat Mater 11, 642–649 (2012). 10.1038/nmat3339

24 Cao, J. et al. Polarized actin and VE-cadherin dynamics regulate junctional remodelling and cell migration during sprouting angiogenesis. Nat Commun 8, 2210 (2017). 10.1038/s41467-017-02373-8

25 Rousseau, S. et al. Vascular endothelial growth factor (VEGF)-driven actin-based motility is mediated by VEGFR2 and requires concerted activation of stress-activated protein kinase 2 (SAPK2/p38) and geldanamycin-sensitive phosphorylation of focal adhesion kinase. J Biol Chem 275, 10661–10672 (2000). 10.1074/jbc.275.14.10661

26 Ferrara, N., Gerber, H. P. & LeCouter, J. The biology of VEGF and its receptors. Nat Med 9, 669–676 (2003). 10.1038/nm0603-669

27 Anderson, S. M., Chen, T. T., Iruela-Arispe, M. L. & Segura, T. The phosphorylation of vascular endothelial growth factor receptor-2 (VEGFR-2) by engineered surfaces with electrostatically or covalently immobilized VEGF. Biomaterials 30, 4618–4628 (2009). 10.1016/j.biomaterials.2009.05.030

28 Anderson, S. M. et al. VEGF internalization is not required for VEGFR-2 phosphorylation in bioengineered surfaces with covalently linked VEGF. Integr Biol (Camb) 3, 887–896 (2011). 10.1039/c1ib00037c

29 Fourel, L. et al. beta3 integrin-mediated spreading induced by matrix-bound BMP-2 controls Smad signaling in a stiffness-independent manner. J Cell Biol 212, 693–706 (2016). 10.1083/jcb.201508018

30 Posa, F. et al. Surface Co-presentation of BMP-2 and integrin selective ligands at the nanoscale favors alpha(5)beta(1) integrin-mediated adhesion. Biomaterials 267, 120484 (2021). 10.1016/j.biomaterials.2020.120484

31 Attwood, S. J. et al. Adhesive ligand tether length affects the size and length of focal adhesions and influences cell spreading and attachment. Sci Rep 6, 34334 (2016). 10.1038/srep34334

32 Hern, D. L. & Hubbell, J. A. Incorporation of adhesion peptides into nonadhesive hydrogels useful for tissue resurfacing. Journal of Biomedical Materials Research 39, 266–276 (1998). 10.1002/(sici)1097-4636(199802)39:2<266::Aid-jbm14>3.0.Co;2-b

33 Rocha, S. F. et al. Esm1 modulates endothelial tip cell behavior and vascular permeability by enhancing VEGF bioavailability. Circ Res 115, 581–590 (2014). 10.1161/CIRCRESAHA.115.304718

34 Cochran, S., Li, C. P. & Ferro, V. A surface plasmon resonance-based solution affinity assay for heparan sulfate-binding proteins. Glycoconj J 26, 577–587 (2009). 10.1007/s10719-008-9210-0

35 Elf, J., Li, G. W. & Xie, X. S. Probing transcription factor dynamics at the single-molecule level in a living cell. Science 316, 1191–1194 (2007). 10.1126/science.1141967

36 Blainey, P. C., van Oijen, A. M., Banerjee, A., Verdine, G. L. & Xie, X. S. A base-excision DNA-repair protein finds intrahelical lesion bases by fast sliding in contact with DNA. Proc Natl Acad Sci U S A 103, 5752–5757 (2006). 10.1073/pnas.0509723103

37 Froese, J. et al. Evolution of SARS-CoV-2 spike trimers towards optimized heparan sulfate cross-linking and inter-chain mobility. Sci Rep 14, 32174 (2024). 10.1038/s41598-024-84276-5

38 Martino, M. M. et al. Extracellular matrix and growth factor engineering for controlled angiogenesis in regenerative medicine. Front Bioeng Biotechnol 3, 45 (2015). 10.3389/fbioe.2015.00045

39 Chandra, P. & Atala, A. Engineering blood vessels and vascularized tissues: technology trends and potential clinical applications. Clin Sci (Lond) 133, 1115–1135 (2019). 10.1042/CS20180155

40 Novosel, E. C., Kleinhans, C. & Kluger, P. J. Vascularization is the key challenge in tissue engineering. Adv Drug Deliv Rev 63, 300–311 (2011). 10.1016/j.addr.2011.03.004

41 Richardson, T. P., Peters, M. C., Ennett, A. B. & Mooney, D. J. Polymeric system for dual growth factor delivery. Nat Biotechnol 19, 1029–1034 (2001). 10.1038/nbt1101-1029

42 Zisch, A. H. et al. Cell-demanded release of VEGF from synthetic, biointeractive cell ingrowth matrices for vascularized tissue growth. FASEB J 17, 2260–2262 (2003). 10.1096/fj.02-1041fje

43 Phelps, E. A., Landazuri, N., Thule, P. M., Taylor, W. R. & Garcia, A. J. Bioartificial matrices for therapeutic vascularization. Proc Natl Acad Sci U S A 107, 3323–3328 (2010). 10.1073/pnas.0905447107

44 Cox, T. R. & Erler, J. T. Remodeling and homeostasis of the extracellular matrix: implications for fibrotic diseases and cancer. Dis Model Mech 4, 165–178 (2011). 10.1242/dmm.004077

45 Trapani, G., Weiss, M. S. & Trappmann, B. Tunable Synthetic Hydrogels to Study Angiogenic Sprouting. Curr Protoc 3, e859 (2023). 10.1002/cpz1.859

46 Farndale, R. W., Buttle, D. J. & Barrett, A. J. Improved quantitation and discrimination of sulphated glycosaminoglycans by use of dimethylmethylene blue. Biochim Biophys Acta 883, 173–177 (1986). 10.1016/0304-4165(86)90306-5

47 Nguyen, D. H. et al. Biomimetic model to reconstitute angiogenic sprouting morphogenesis in vitro. Proc Natl Acad Sci U S A 110, 6712–6717 (2013). 10.1073/pnas.1221526110

48 Thakar, D. et al. A quartz crystal microbalance method to study the terminal functionalization of glycosaminoglycans. Chem Commun (Camb) 50, 15148–15151 (2014). 10.1039/c4cc06905f

49 Bartelt, S. M. et al. Dynamic blue light-switchable protein patterns on giant unilamellar vesicles. Chem Commun (Camb) 54, 948–951 (2018). 10.1039/c7cc08758f

50 Di Iorio, D., Lu, Y., Meulman, J. & Huskens, J. Recruitment of receptors at supported lipid bilayers promoted by the multivalent binding of ligand-modified unilamellar vesicles. Chem Sci 11, 3307–3315 (2020). 10.1039/d0sc00518e

51 Lind, T. K. & Cardenas, M. Understanding the formation of supported lipid bilayers via vesicle fusion-A case that exemplifies the need for the complementary method approach (Review). Biointerphases 11, 020801 (2016). 10.1116/1.4944830

52 Di Iorio, D. & Wegner, S. V. Dynamic Light-Induced Protein Patterns at Model Membranes. J Vis Exp (2024). 10.3791/66531

