## Supporting Information for "Heparin flexibility within the extracellular matrix determines the bioactivity of bound vascular endothelial growth factor"

* Correspondence

Prof. Dr. Britta Trappmann

| Peptide type | Peptide sequence | Net charge at pH 7.0 |
|---|---|---|
| Crosslinker | **K**C**D**GVPMSM**R**GGC**K** | +1.91 |
| Mono-Cys peptide | **K**C**D**GVPMSM**R**GGG**K** | +1.91 |

**Supplementary Table 1. Amino acid sequence and net charge of synthetic hydrogel crosslinker and mono-Cys peptide.** Single-letter amino acid sequence of peptides: positively charged amino acids are labeled in red, negatively charged amino acids in blue, underlined letters indicate sites for coupling onto biopolymer chains. Net charge at pH 7.0 was calculated using the openly available peptide and amino acid calculator provided online by Bachem (https://www.bachem.com/knowledge-center/peptide-calculator/).

| DexHepMA hydrogel composition | | Relative matrix positive charge density |
|---|---|---|
| Crosslinker | Mono-Cys peptide | |
| 17.4 mM | - | 0.7 |
| 17.4 mM | 25.9 mM | 1.7 |
| 26 mM | - | 1 |
| 26 mM | 17.3 mM | 1.7 |
| 43.4 mM | - | 1.7 |

**Supplementary Table 2. Composition of hydrogels with varying positive matrix charge density.** In our approach, differences in the matrix positive charge density only result from changes in crosslinker and mono-Cys peptide concentrations (since all other hydrogel ingredients, such as DexMA, HepMA or the adhesive peptide CGRGDS remain constant). Hence, to facilitate comparison between hydrogels, the positive charge density of each hydrogel is reported relative to the positive charge density of a DexHepMA hydrogel crosslinked with 26 mM crosslinker peptide (in absence of mono-Cys peptide), while taking into account net charges of the crosslinker and mono-Cys peptide and their concentrations only.

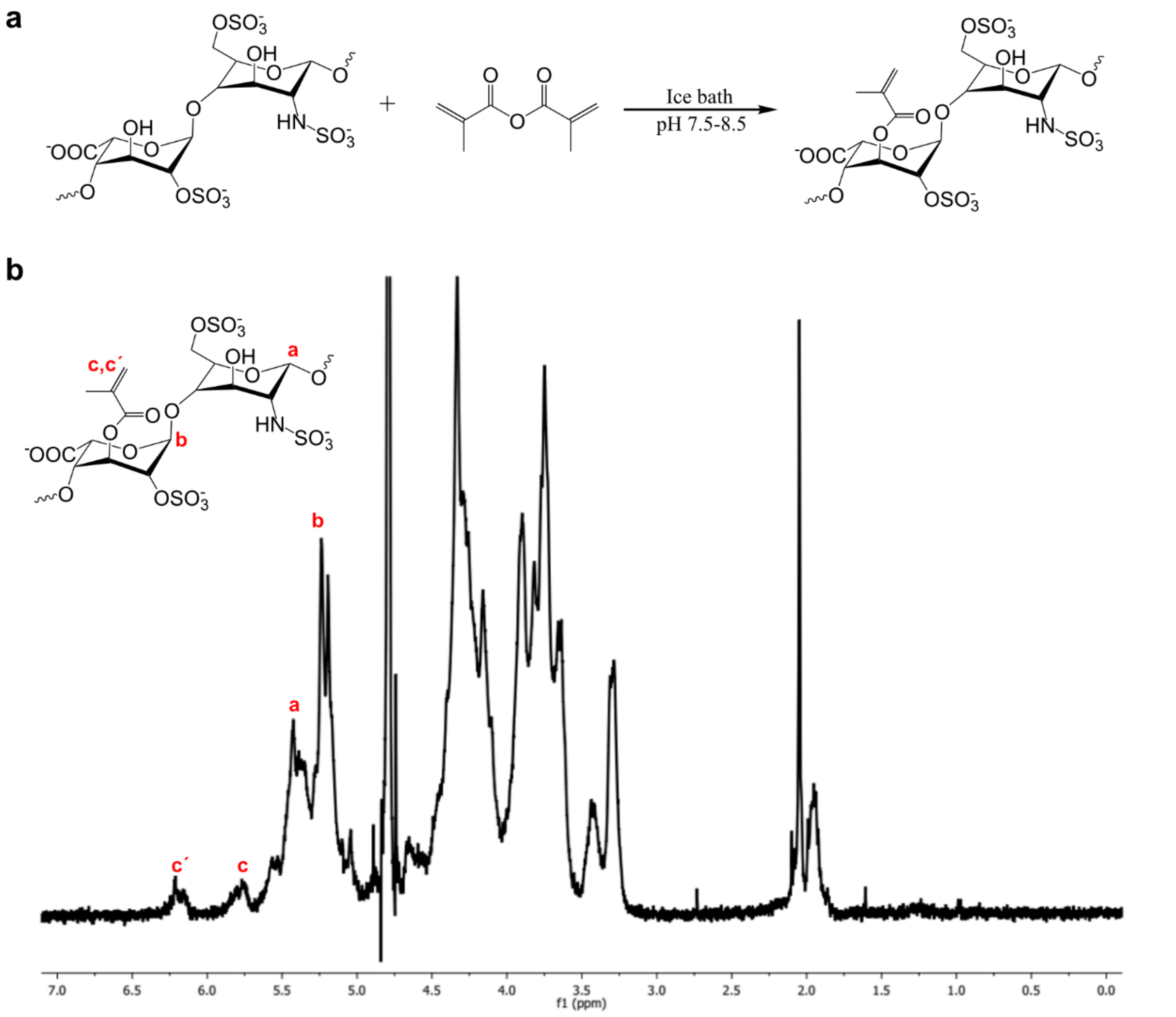


**Supplementary Figure 1. Methacrylation of heparin. a** Heparin is reacted with methacrylic anhydride to obtain methacrylated heparin (HepMA). **b** $^{1}$H-NMR spectrum of HepMA. The structure of the heparin major disaccharide unit is reported for peak assignment.

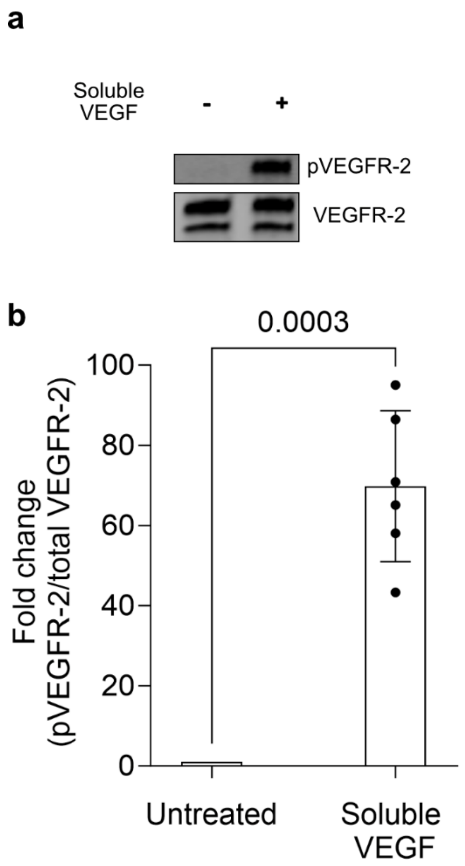


**Supplementary Figure 2. HUVECs are responsive to soluble VEGF. a** HUVEC monolayers were exposed to soluble VEGF (30 ng/mL) or EBM-2 medium only for 2 min, followed by immunoprecipitation of VEGFR-2 and Western blot analysis for pan-phosphorylation of tyrosine residues (using 4G10 antibody) (top panel). As loading control, the blots were reprobed with an antibody against total human VEGFR-2 (bottom panel). **b** Quantification of fold change of pVEGFR-2 signal intensities, adjusted to the level of precipitated VEGFR-2 and normalized to the untreated sample. All data are reported as mean ± SD, $p < 0.05$ is considered to be statistically significant (two-tailed unpaired t-test with Welch´s correction). (n = 6 independent samples).

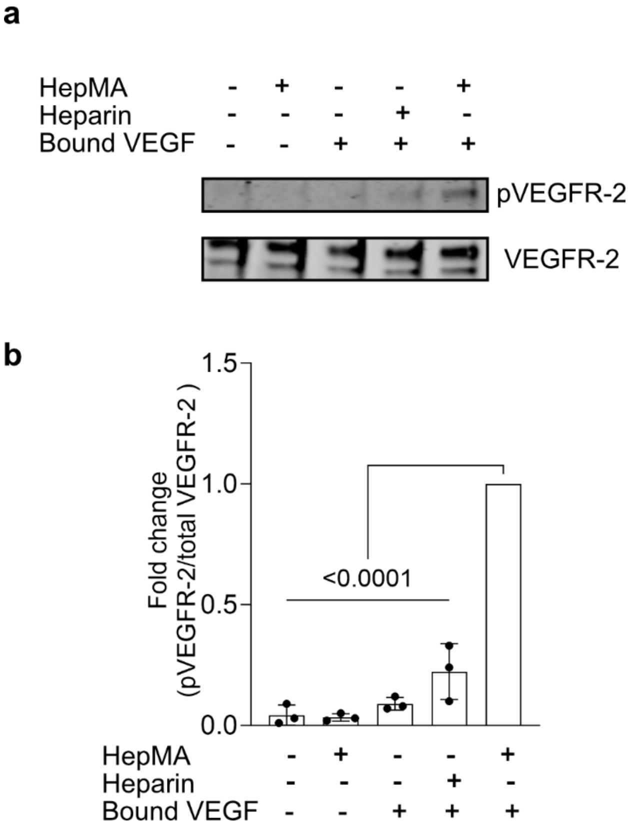


**Supplementary Figure 3. VEGFR-2 activation only occurs when VEGF is bound to hydrogels containing methacrylated heparin. a** HUVEC monolayers were exposed for 2 min to different formulations of dextran-based hydrogels crosslinked with 26 mM crosslinker, with (+) /without (-) either HepMA or heparin, and with (+) /without (-) matrix-bound VEGF (330 ng/mL), followed by immunoprecipitation of VEGFR-2 and Western blot analysis for pan-phosphorylation of tyrosine residues (using 4G10 antibody) (top panel). As loading control, the blots were reprobed with an antibody against total human VEGFR-2 (bottom panel). **b** Quantification of fold change of pVEGFR-2 signal intensities, adjusted to the level of precipitated VEGFR-2 and normalized to the HUVEC sample treated with a VEGF-functionalized (330 ng/mL) DexHepMA hydrogel containing 26 mM crosslinker. All data are reported as mean ± SD, $p < 0.05$ is considered to be statistically significant (one-way ANOVA with Dunnett´s post-hoc test). ($n = 3$ independent samples).

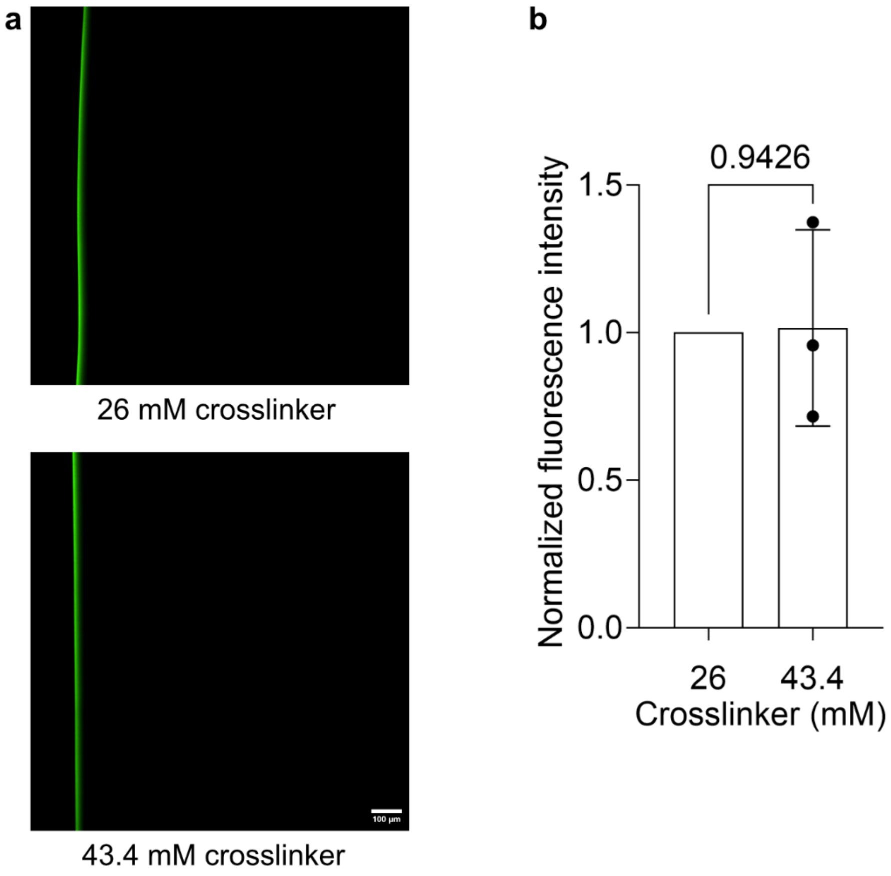


**Supplementary Figure 4. Concentration of VEGF bound to synthetic hydrogels is independent of crosslinker concentration. a** FITC-labeled VEGF was added to the source channel of a microfluidic device[47] and allowed to bind to DexHepMA hydrogels of varying crosslinker concentration for 2 h. Bound FITC-labeled VEGF was imaged following overnight washes in PBS. Scale bar, 100 µm. **b** Fluorescence intensity of channel surface-bound VEGF was measured as the highest intensity value of the plot profile acquired perpendicular to the channel wall. Fluorescence intensity values were averaged from 25 measurements per image ($n = 3$ independent samples) and normalized to a DexHepMA hydrogel sample containing 26 mM crosslinker. All data are reported as mean ± SD, $p < 0.05$ is considered to be statistically significant (two-tailed unpaired t-test with Welch´s correction).

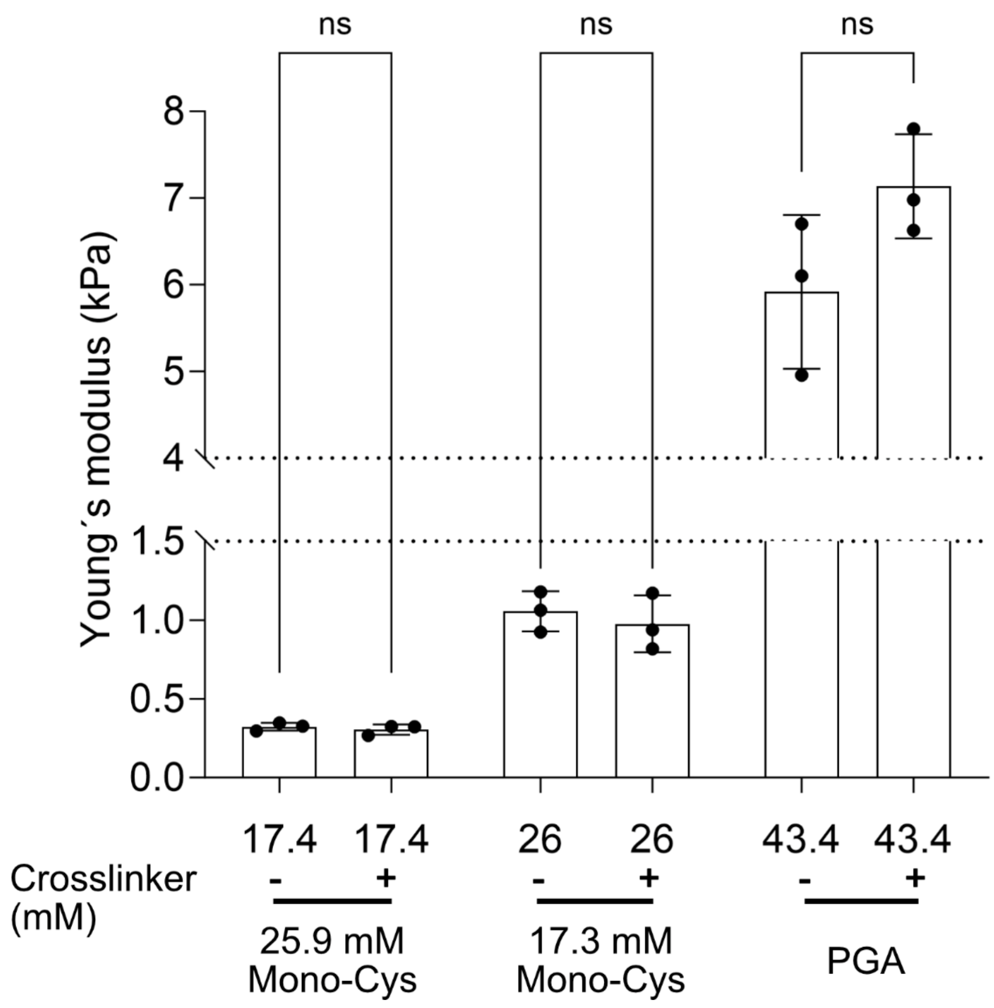


**Supplementary Figure 5. Mechanical characterization of synthetic hydrogels.** Young's moduli of DexHepMA hydrogels with varying concentrations of crosslinker and composition: 17.4 mM crosslinker either with (+) or without (-) mono-Cys peptide (25.9 mM), 26 mM crosslinker either with (+) or without (-) mono-Cys peptide (17.3 mM), 43.4 mM crosslinker either with (+) or without (-) PGA. All data are reported as mean ± SD. For pairwise comparisons two-tailed unpaired Student's t-test was used. Exact p values: (from left to right) ns = 0.4860, ns = 0.5678, ns = 0.1210. (n = 3 independent samples).

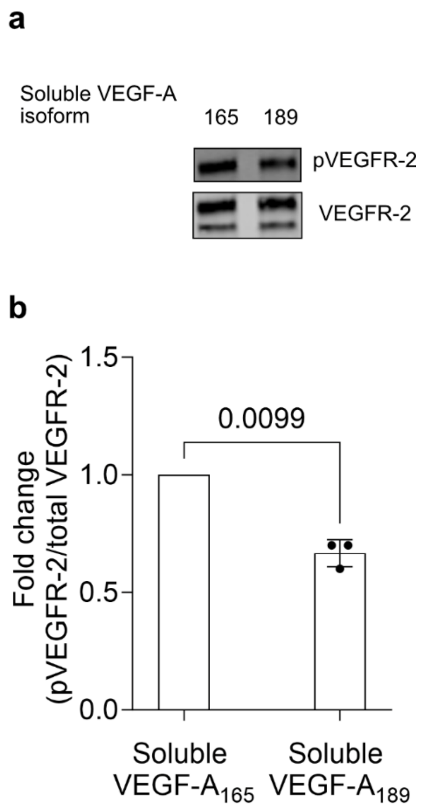


**Supplementary Figure 6. Bioactivity of soluble VEGF-A189. a** HUVEC monolayers were exposed to either soluble VEGF-$A_{165}$ (30 ng/mL) or soluble VEGF-$A_{189}$ (30 ng/mL) for 2 min, followed by immunoprecipitation of VEGFR-2 and Western blot analysis for pan-phosphorylation of tyrosine residues (using 4G10 antibody) (top panel). As loading control, the blots were reprobed with an antibody against total human VEGFR-2 (bottom panel). **b** Quantification of fold change of pVEGFR-2 signal intensities, adjusted to the level of precipitated VEGFR-2 and normalized to the sample treated with soluble VEGF-$A_{165}$. All data are reported as mean ± SD, $p < 0.05$ is considered to be statistically significant (two-tailed unpaired t-test with Welch´s correction). (n = 3 independent samples).

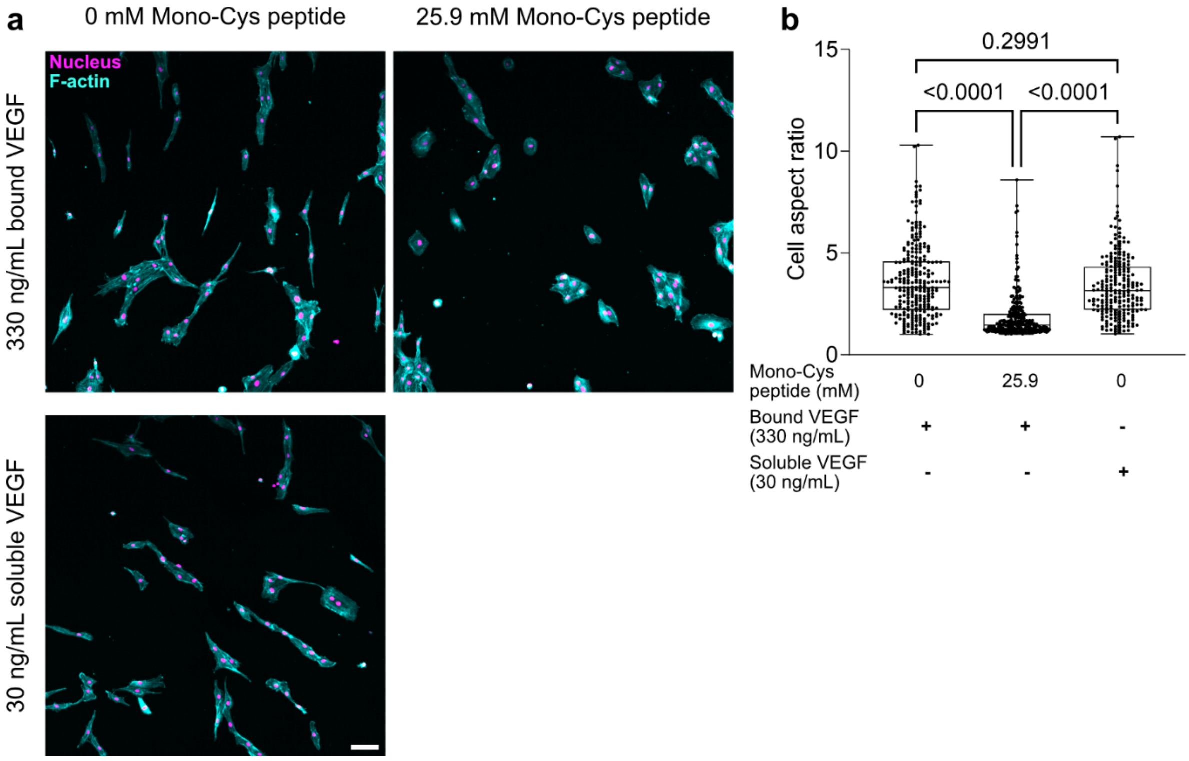


**Supplementary Figure 7. Mobility of matrix-bound VEGF regulates HUVEC elongation.** **a** Composite fluorescence images show nuclei (magenta) and F-actin (cyan) of HUVECs cultured overnight (>16 h) atop DexHepMA hydrogels, with (right) or without (left) 25.9 mM mono-Cys peptide, crosslinked with 17.4 mM crosslinker peptide in presence of either bound (330 ng/mL) or soluble (30 ng/mL) VEGF. **b** Quantification of cell aspect ratio for conditions shown in (**a**) (n > 200 cells, pooled from 3 independent experiments). Data are reported as box-and-whiskers plots (box, 25–75 percentile; bar-in-box, median; whiskers, the minimum and maximum values) (one-way ANOVA with Tukey´s post-hoc test). Scale bar, 100 µm.

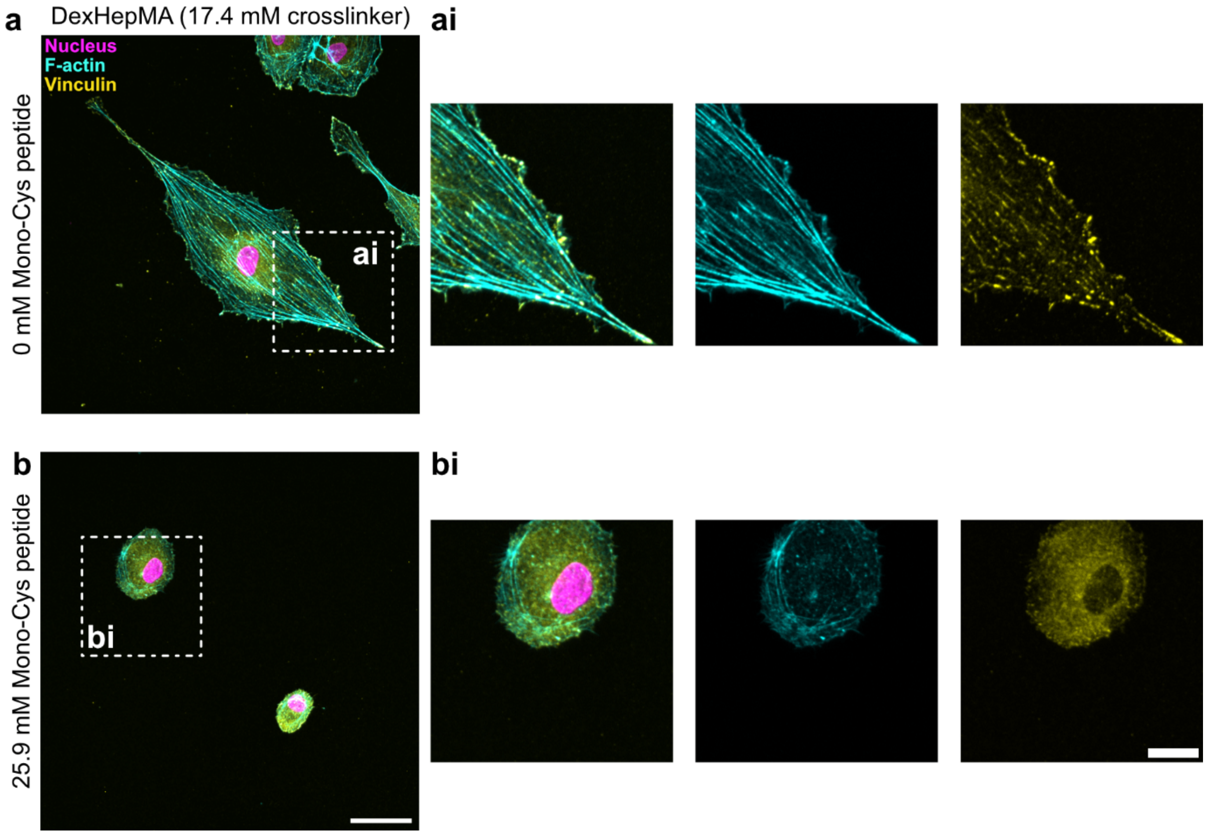


**Supplementary Figure 8. Mobility of matrix-bound VEGF regulates focal adhesion assembly.** High magnification composite fluorescence images show nuclei (magenta), F-actin (cyan) and vinculin (yellow) of HUVECs cultured overnight (>16 h) atop DexHepMA hydrogels crosslinked with 17.4 mM crosslinker in absence of mono-Cys peptide (**a**, **ai**) or with 25.9 mM mono-Cys peptide (**b**, **bi**) and functionalized with 330 ng/mL VEGF. Scale bar: 100 µm, scale bar in inset: 20 µm.
